# Controlling Flow Dynamics and Permeability in Perfusable Vascular Constructs Using Volumetric 3D Printing

**DOI:** 10.1101/2025.08.11.669590

**Authors:** Julia Simińska-Stanny, Agathe Thiry, Ilargi Balda Lorenzo, Adam Junka, Kacper Pietrzak, Marco De Corato, Maria Jose Gomez-Benito, Armin Shavandi

## Abstract

Creating perfusable vascular networks that replicate physiological flow remains a key challenge in tissue engineering. Here, we present a volumetric 3D printing (Vol3DP) method for fabricating tunable, biologically relevant vascular structures that enable controlled perfusion. A recyclable resin of methacrylated gelatin (GelMA) and polyethylene glycol diacrylate (PEGDA) was optimized for Vol3DP to produce high-fidelity hydrogels with embedded channels. To evaluate the relationships between flow, structure, and function, we designed modular perfusion platforms that offer precise control over physiological shear stress (3–50 dyne/cm²), flow rates (1–15 mL/min), and flow modes, including pulsatile and continuous. These platforms support endothelial cell attachment and spreading under static seeding conditions, sustained perfusion, and permeability assessment and further allow for direct comparisons of flow dynamics and perfusion efficiency in channel-in-hydrogel systems. The finally engineered adaptable platform allowed for independent pressure modulation within the vessel and the outside environment. We conducted simulations of flow and hydrogel permeability that closely resemble experimental results. These results underscore the dominance of diffusive transport through the hydrogel matrix and highlight the ability of our system to simulate physiologically relevant mass transfer phenomena. Finally, we conducted a pilot study using *Galleria mellonella* larvae to compare active and passive dye transport *in vivo*, complementing *in vitro* data and highlighting the importance of perfusion-capable scaffolds for accurately simulating vascular drug delivery environments.

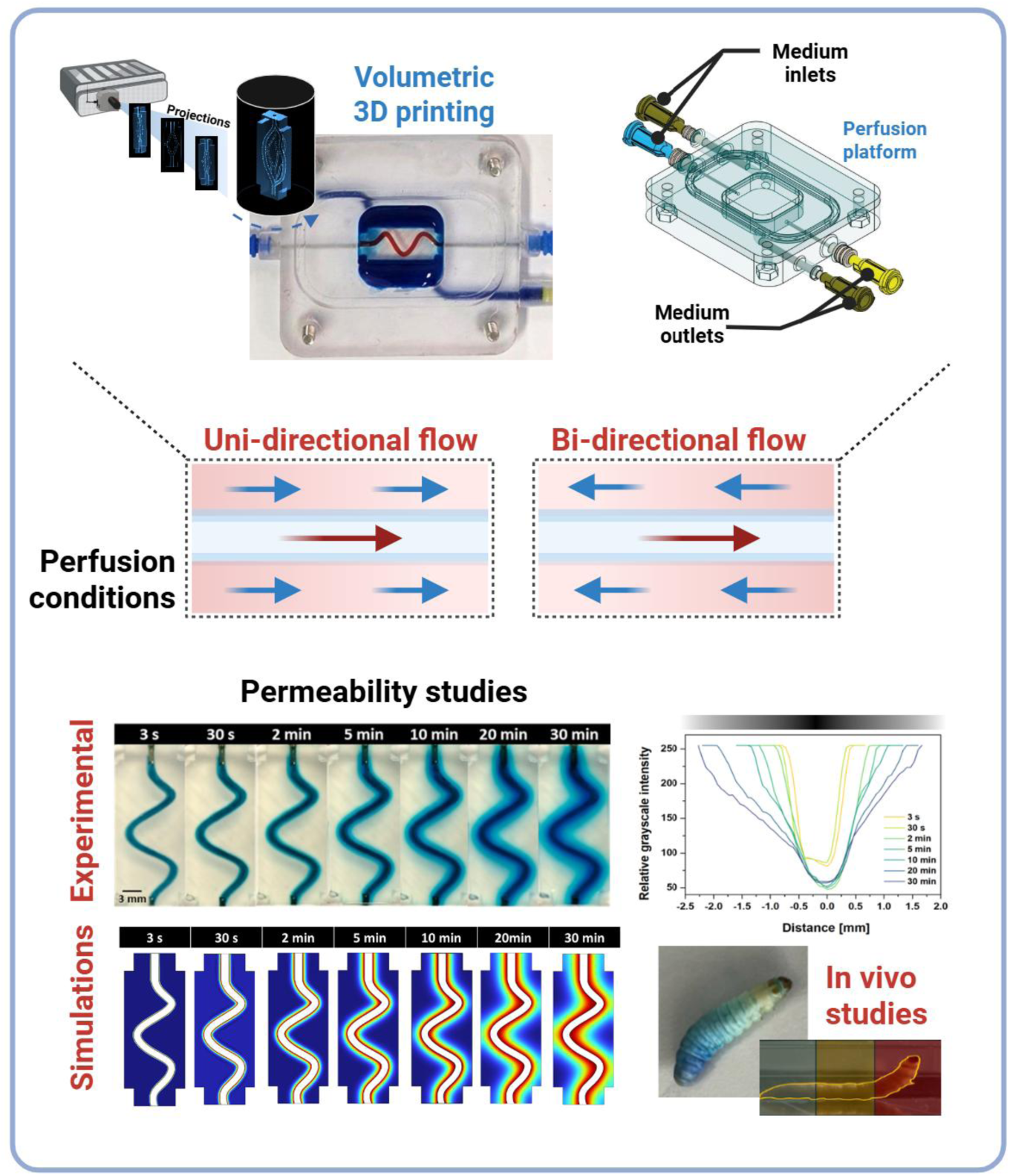

## 1. Introduction

Biofabrication techniques have transformed tissue engineering by enabling precise control over the 3D arrangement of cells and biomaterials [1]. However, recreating vascular networks remains a crucial aspect of successful tissue engineering, and it remains a significant challenge [2, 3].

Native blood vessels exhibit a hierarchical organization from large arteries and veins to microcapillaries. Their walls consist of three layers: the tunica intima (endothelial monolayer supported by basement membrane), tunica media (smooth muscle cells and elastic lamellae), and tunica adventitia (fibroblasts, collagen fibers, and vasa vasorum). The ECM is predominantly composed of type I and III collagen, elastin, proteoglycans, and glycoproteins, providing both mechanical strength and bioactive cues for vascular homeostasis [4].

Static culture conditions are insufficient to recapitulate the dynamic environment of living tissues [5]. To date, most perfusable vascular models are confined to microfluidic channels [6], which are commonly categorized as organ-on-chip platforms [7, 8]. These systems are primarily fabricated using lithography techniques, including polydimethylsiloxane (PDMS) molding, to create channels that serve as substrates for cell culture. While these platforms approximate some physiological dynamics, their reliance on rigid structures and small geometries limits their broader applicability in tissue engineering. Hydrogels are increasingly employed in vascular models, providing a physiologically relevant 3D environment that supports the seeding of endothelial cells (ECs) and the formation of vascular networks through vasculogenesis [7, 9]. Although these networks enable the study of barrier function, their randomly oriented geometries limit flow studies or define wall shear stresses [7, 10]. In some cases, channels are created within PDMS and then coated or shaped within hydrogels [11, 12]. For example, vascular geometries can be fabricated by casting a hydrogel around a steel needle, where the lumen is formed upon removal of the needle. However, this approach is limited to creating straight channel geometries [11] with the channel size determined by the needle diameter, typically 60-300 μm [13, 14]. Although organ-on-chip platforms aim to replicate native dynamic conditions, they often require multiple components and involve a cumbersome assembly process [15]. To address the limitations of existing vascular models, researchers use 3D printing techniques [16]. Significant efforts have been made to fabricate vascular structures using extrusion bioprinting [17], sacrificial embedded printing [18], melt electrowriting [19], digital light processing (DLP) [20, 21], and hybrid approaches [16, 22, 23]. While these methods have achieved channeled geometries, their lengthy fabrication time and difficulties in assembling fluid-circulating setups remain major drawbacks. Volumetric printing (Vol3DP) is a promising approach to vascular printing that can rapidly sculpt photoresponsive hydrogels into complex 3D structures, including channeled geometries [24], within approximately one minute [1, 25].

Unlike extrusion bioprinting, which typically produces features >100 µm in diameter, Vol3DP can reproducibly fabricate features down to ∼40–50 µm, enabling the recreation of microvascular geometries with higher anatomical fidelity. Using principles inspired by computed tomography (CT), Vol3DP employs cumulative light projections from multiple angles to crosslink bioresins, forming the entire structure simultaneously when the energy dose surpasses the photocrosslinking threshold [1, 25]. Preferred materials for Vol3DP are high-viscosity (∼90,000 cP) or solid, thermally gelled resins, which minimize relative motion between the printed object and the uncured resin, thereby reducing molecular diffusion–induced blurring [25, 26]. Among Vol3DP resins for bio applications, gelatin has emerged as a leading material, due to its favorable thermal gelation and ease of modification [23, 27, 28]. However, gelatin alone often fails to produce mechanically robust constructs capable of maintaining their shape [26]. Polyethylene glycol diacrylate (PEGDA), though limited by its low viscosity (∼120 cP), complements gelatin by enhancing mechanical stability when used in combination [29].

Whilst Vol3DP solves numerous difficulties of current vascular tissue biofabrication, a substantial portion of research within this field focuses on material explorations and printing process refinement. Successful fabrication of channel structures has been demonstrated, but the development of systems with highly controllable perfusion remains underexplored [1, 19, 30]. Using flow platforms to dynamize cell cultures within hydrogels can provide insights into how ECs sense molecular cues and regulate cell-cell adhesions, modulating gas, ion, and other transports [31, 32]. Additionally, such platforms can be utilized in the future to study flow-driven morphological changes in EC alignment, enhancements of adhesion bonds, cell-cell junctions, or cytoskeletal developments, using appropriate immunofluorescent staining [11, 33].

From a model validity standpoint, within the same vessel, shear stress levels and flow profiles can vary significantly due to geometric features, including vessel branches and curvature [34]. Recapitulating these particularities has not been possible using wire-templating or extrusion printing techniques.

Alterations in shear stress are implicated in the onset and progression of vascular diseases, including atherosclerosis, aneurysms, and thrombosis [35]. Models capable of reproducing physiological shear stress profiles are therefore essential for studying mechanobiological disease mechanisms and evaluating therapeutic strategies.

In this work, we present a novel strategy to introduce controlled flow dynamics, as previously achieved for microfluidic PDMS vascular models, but this time integrating flow constraints with structures fabricated through Vol3DP methods. Previous studies have employed GelMA–PEGDA in Vol3DP [36], demonstrating high print fidelity but without optimization for resin reusability or integration with modular perfusion platforms. In contrast, our work systematically optimizes optical/mechanical balance, evaluates photoinitiator toxicity trade-offs, and introduces reusability testing for sustainability. To replicate physiological conditions, we develop an *in vitro* platform for dynamic perfusion.

Although the presented resin formulation offers printability, mechanical stability, and optical clarity, it remains compositionally distinct from native vascular ECM, which is rich in collagen and elastin fibers. Future work will focus on integrating ECM-derived biopolymers to more closely mimic the biochemical cues of native tissue while preserving the high-fidelity printing performance of Vol3DP resins. This system is designed to simulate physiological shear stresses and flow rates, supporting a range of fluid-handling configurations, including syringe pumps, peristaltic systems, and rocker platforms. The modular design ensures compatibility with Vol3DP-fabricated vascular models, enabling versatile experimental setups to study tissue engineering applications under diverse flow conditions. The proposed platform offers a valuable tool for investigating vascular biology, drug delivery, and the development of clinically relevant *in vitro* models.

## 2. Results and discussion

### 2.1 Biomaterial resin formulation and volumetric bioprinting

Hydrogels composed of GelMA and PEGDA have been widely used in extrusion-based bioprinting [37, 38]; however, their adaptation to Vol3DP requires optimization. In multi-component systems, differences in optical clarity, viscosity, and polymerization kinetics between components can lead to uneven crosslinking or loss of resolution during volumetric exposure. While several studies have successfully printed multi-material resins via Vol3DP [30, 39], each formulation requires tailored adjustments of printing parameters to accommodate the specific physicochemical interactions between its components.

In Vol3DP, thermally solidified GelMA can hold the print through multiple rotation sequences when 2D images are projected to cure the material selectively [40] (**Figure 1A**). GelMA was selected as the base material due to its biocompatibility, enzyme-mediated biodegradability, and thermo-reversible properties, which facilitate structural integrity during bioprinting [24]. Polyethylene glycol diacrylate (PEGDA) was added to enhance mechanical stability and print fidelity [26]. Nine resin formulations were prepared, varying GelMA (5 %, 10 % w/v) and PEGDA (5 %, 10 % w/v) concentrations, along with lithium phenyl(2,4,6-trimethylbenzoyl)phosphinate (LAP) photoinitiator (0.3, 0.5, 1.0 mg/mL), (**Supplementary information (SI), Table S1 –** Formulation of biomaterial resins), based on the previous studies exploring GelMA printability [23, 27, 28].

**Figure 1.**
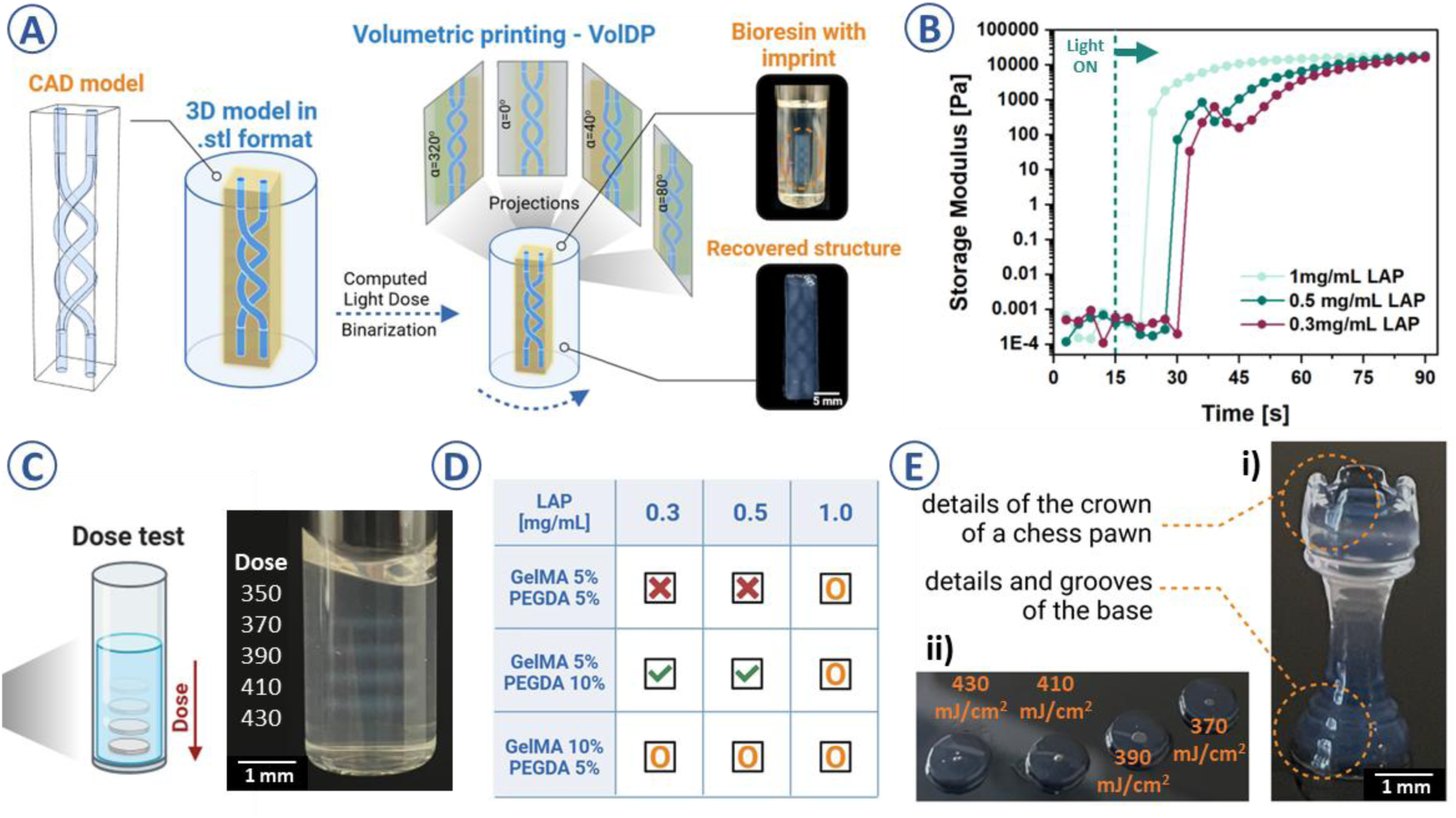
**A)** Schematic of Vol3DP of double-spiral channeled geometry, with the imprinted geometry and the recovered construct; **B)** Time-sweep test showing the photocrosslinking pattern for variously formulated bioresins, the dashed line represents the moment from which the 405 nm light was on, representative plots, n=3; **C)** Schematic of the light dose optimization for resins at different light doses, performed dose test is used to determine the light exposition of the material required to photocrosslink the projected patterns with an example of the test used to adjust the dose for GelMA 5 % PEGDA 10 % biomaterial resin; **D)** Volumetric printability results for various bioresins with different LAP content (derived based on the results from Figure S1). The green tick indicates high-detail printability, the orange circle denotes printability with compromised detail and reduced stability, and the red ‘x’ marks poorly printable samples, in which the disc structures were not distinguishable. **E)** Vol3DP results for GelMA 5 % PEGDA 10 % LAP 0.3 mg/mL biomaterial resin (i) disks printed with different light doses and (ii) chess figure revealing the structural details of the print.

The optimal light dose was selected based on quantitative metrics, including the dimensional accuracy of disc-shaped structures [28, 41] measured via coherence-contrast microscopy and benchmark geometries such as a chess figure (**Figure 1C-E**, **SI, Figure S1**). Based on a comparative evaluation, we identified 370 mJ/cm² as the optimal dose for the GelMA 5% + PEGDA 10% + LAP 0.3 mg/mL formulation. This dose provided the best balance between print fidelity and mechanical robustness. At 350 mJ/cm², the constructs exhibited insufficient curing, resulting in deformation and poor feature resolution. At 400 mJ/cm², over-polymerization effects, such as blurred features and reduced translucency, were observed. The 370 mJ/cm² dose produced stable, self-standing structures with a resolved geometry, as confirmed by coherence-contrast microscopy and photorheology.

The results indicated that increasing the PEGDA concentration improved structural stability and resolution [43], as demonstrated by constructs printed with 10% PEGDA and 5% GelMA, however increasing PEGDA content shifts the resin composition further from native ECM biochemistry, a limitation to be addressed in future ECM-enriched formulations.

Formulations containing higher GelMA concentrations (10%) resulted in increased resin turbidity, likely due to enhanced molecular aggregation and light scattering, which reduced printing resolution and led to fragile, difficult-to-retrieve objects. Conversely, lowering GelMA to 5% while increasing PEGDA to 10% improved structural stability and transparency (**SI, Figure S1**). We also varied the LAP photoinitiator concentration (0.3–1 mg/mL) to determine the minimum amount required for successful printing while maintaining resolution and potentially reducing cytotoxicity. Photorheology tests showed that the formulation with lower LAP concentration (0.3 mg/mL) had a slower, controlled polymerization kinetics (peak modulus reached in ∼30 s), allowing gradual curing that minimized overcuring and increased printing accuracy.

In contrast, for our specific formulation and standard dose settings, higher LAP concentrations (1 mg/mL) resulted in faster polymerization (∼10 s), which, without dose reoptimization, led to reduced feature definition and signs of overcuring (**Figure S1**). Other studies have successfully employed 1 mg/mL LAP in volumetric printing [23, 30]. Our observations should thus be interpreted as formulation and dose-specific, rather than as a general limitation of high LAP concentrations. Photorheology (**Figure 1B**) further indicates the importance of controlled polymerization, especially in chain growth systems such as GelMA and PEGDA [24, 27]. Following the dose experiments, The 5% GelMA + 10% PEGDA + 0.3 mg/mL LAP combination was selected for its balance of mechanical stability, optical clarity, and reduced LAP content to improve cell compatibility.

### 2.2 Biomaterial-resin Transparency and Print Resolution

The effectiveness of Vol3DP relies on materials with high optical transparency and minimal light scattering or absorption [42, 43]. To crosslink the resin, light must pass through the entire build volume without distortion or attenuation, making resin transparency and the photoinitiator concentration crucial [44].

We compared our resin to GelMA (10% with 1 mg/mL LAP), which is commonly used in Vol3DP. The resin had higher clarity than GelMA resin when loaded into printing vials (**Figure 2A**). Prints of a channeled object showed uneven edges and over-polymerization in GelMA, while resin produced precise striations, typical of Vol3DP, confirming its precision curing (**Figure 2B**).

**Figure 2.**
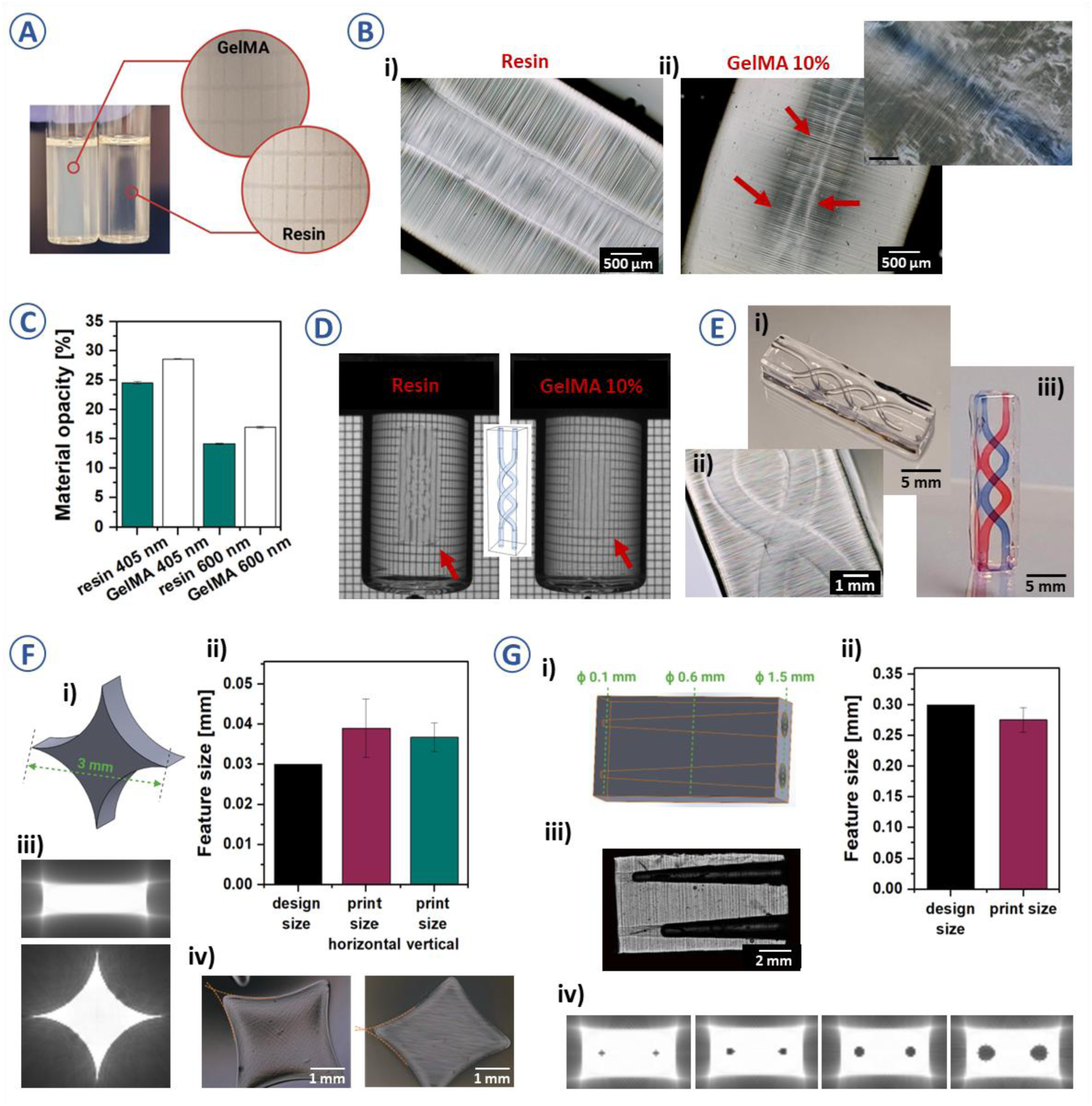
**A)** Photographs of the GelMA and resins when loaded to the printing vials, the grid behind the vial is less visible for GelMA resin; **B)** Vol3DP of resin showed sharper and clearer internal channels compared to GelMA; **C)** Material opacity results obtained based on absorbance measurements for GelMA and both resin and hydrogel at 405 nm and 600 nm, n=4, mean ± SD., differences between groups were significant at p ≤ 0.001, two-way ANOVA with Tukey post-hoc test; **D)** Images of the vials after Vol3DP of double-channeled structure, the object is better visible for resin than for GelMA likely due to a higher differences in refractive indexes between the resin and hydrogel formed due to photocrosslinking; **E)** Images of the Vol3DP double-channel structure, emphasizing on the structural details (ii) and perfusability of the channels (i, iii); **F)** Vol3DP results of the highest-resolution positive features (i) shows the designed model, (ii) comparison of the printed feature size with the model size, n=3, mean ± SD., data were not significant at p = 0.05, Kruskal-Wallis with Dunn-Sidak post-hoc test; (iii) software-generated light projection patterns, corresponding to the design, (iv) Vol3DP structures used for evaluation **G)** Vol3DP results of the highest-resolution negative features (i) shows the designed model, (ii) comparison of the printed feature size with the model size, n=3, mean ± SD., data were not significant at p = 0.05, Kruskal-Wallis with Dunn-Sidak post-hoc test; (iii) Vol3DP structures used for evaluation, (iv) software-generated light projection patterns, corresponding to the design.

Lower absorbance values of the resin at 405 nm (24.17%) compared to pure GelMA (28.60%) indicate reduced attenuation of curing light, directly translating to higher curing precision for the resin. Measurements at 600 nm, although not directly related to the curing wavelength, support the overall superior transparency and lower optical scattering within the resin, thereby enhancing overall print precision (**Figure 2C**). A greater refractive index contrast was observed between the uncured resin and its corresponding hydrogel (∼0.06%) compared to GelMA, which contributed to the enhanced visibility of the resin-printed structures during the volumetric curing process (**Figure 2D**, **SI, Figure S2-C**). The high turbidity in GelMA and resins with higher GelMA content (**SI, Figure S1)** is likely due to greater heterogeneity within the resin matrix, possibly involving the formation of larger polymer aggregates, resulting in increased light scattering and reduced precision in light-guided curing (**Figure 2D**) [42]. We hypothesize that the compatibility of GelMA and PEGDA for high-resolution multimaterial volumetric printing can be attributed to their similar light doses for polymerization, comparable photocrosslinking patterns (**SI, Figure S2-A**), and closely matched refractive index values prior to crosslinking (**SI, Figure S2-C**).

Although previous studies report that polymerizing 5% (w/v) GelMA with 0.1% (w/v) LAP requires ∼250 mJ/cm² [1], and PEGDA hydrogels require ∼236 ± 14 mJ/cm², these values serve only as reference points. In our resin formulation, the polymerization behavior reflects a new composite system, where the optimal dose depends on the individual polymer thresholds and concentration, molecular weight, degree of functionalization, and the non-orthogonal interaction between GelMA and PEGDA radical polymerizations. Therefore, empirical optimization of the light dose was essential to ensure proper crosslinking, and the final dose (370 mJ/cm²) was selected based on print fidelity and mechanical integrity. Adjusting the light dose and avoiding overexposure, we could print precise structures with embedded perfusable channels (**Figure 2E**).

Coherence contrast microscopic analysis was performed post-3D printing to evaluate the maximum resolution of the Vol3DP models. Two geometries were printed, emphasizing either positive or negative features (**Figures F-i and G-i**). The minimum thickness of quadrangle tapering was 49 ± 7 µm for horizontal prints and 43 ± 4 µm for vertical prints, [1, 45].

The achieved resolution limits (approximately 40–50 µm) closely matched the model dimensions, confirming the high printability and resolution of both the material and Vol3DP method. This resolution is sufficient for modeling intermediate microvascular structures with meaningful biological relevance.

A clear distinction was observed between the printable and non-printable diameters (**Figure 2G, iii and iv**). The design remained successfully printable until the channel diameter reached 186 ± 28 µm, beyond which perfusability was compromised (**Figure 2G-iii**), establishing the resolution limit for negative concave features at ∼200 µm. While this resolution enables the modeling of intermediate-sized vessels, such as arterioles and venules, it is insufficient to replicate physiological capillary networks, which typically range in diameter from 5 to 10 µm. This is a limitation of the current system, particularly for applications that require the accurate recreation of fine-scale vascular beds, such as microphysiological systems or organ-on-chip platforms. Future integration approaches, such as sacrificial templating or cell-guided microvascular assembly, may help address this constraint.

The light intensity patterns simulated using the printer software (Apparite®) enabled predictions of potential over-curing areas and minimal printable diameters (**Figure 2F-iii, G-iv**). This facilitated iterative adjustment to the design constraints to achieve perfusable structures.

### 2.3 Bioresin Stability, Reusability and Rheological Properties

A semi-solid gelatin network can stabilize PEGDA monomers during the Vol3DP printing process, preventing the crosslinked material from settling down during light exposure and rotation. The physical crosslinking of GelMA when cooled to below 25°C enables the formation of hydrogen bonds between the GelMA polymer chains, leading to increased storage modulus (G’) [46, 47]. Similarly high G’ (≥ 100 Pa) were observed for the resin at temperatures <30°C (**Figure 3A-i**). When the temperature reaches ∼30°C for the resin, a sharp drop in G’ occurs, indicating a thermal gel-sol transition. At this point, viscous behavior starts to dominate over elastic behavior (flow point) [48]. The material loses its rigidity, becomes soft (G’ < 0.001 Pa), and begins to flow. The exact sol-gel transition temperature can be influenced by the concentration of GelMA and its degree of methacrylation [49, 50]. This transition is reversible and beneficial for Vol3DP, as it allows easy recovery of the crosslinked object, unaffected by temperature changes when the surrounding material liquefies (**Figure 3A-ii**).

**Figure 3.**
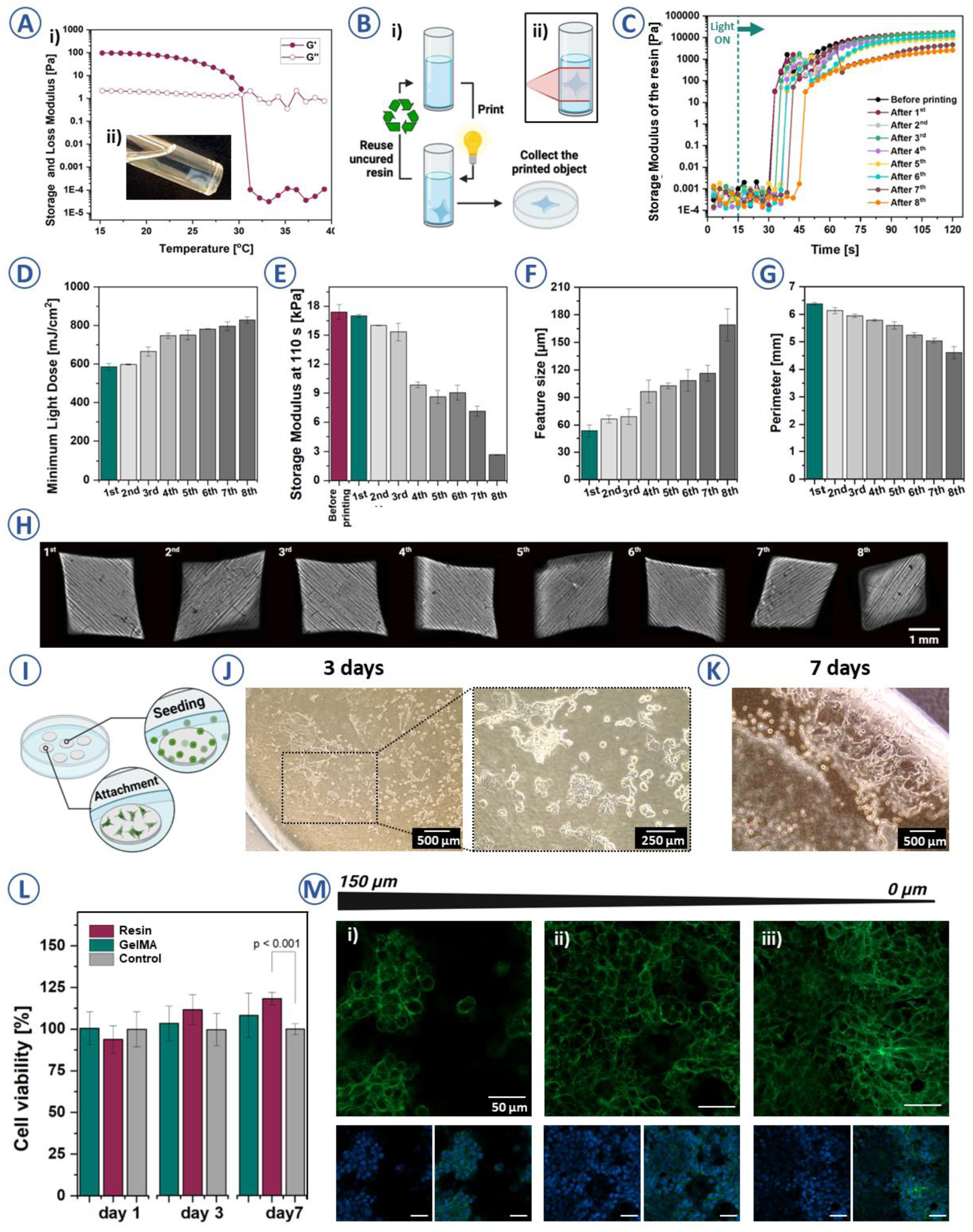
**A)** Temperature sweep test for resin – (i) (representative plot based on n=3) and (ii) Vol3DP shape remaining solid once the rest of the resin is liquified; **B)** Schematic illustration of the resin reusability concept, where the unpolymerized resin is recovered and used for subsequent prints (i), (ii) – exposure of the material volume during object printing; **C)** Photorheology tests for resin after every Vol3DP cycle (representative plots based on n=3), the dashed line represents the moment from which the 405 nm light was on; **D)** Minimum light dose required to crosslink the resin after repeated printing n=3, mean ± SD., results were significant at p ≤ 0.05 for 1^st^ vs. 8^th^ print and 2^nd^ vs. 8^th^ print, Kruskal-Wallis; **E)** Perimeter of the Vol3DP design after consecutive printing, n=3, mean ± SD., results were significant at p ≤ 0.05 only for 1^st^ vs. 8^th^ print, Kruskal-Wallis; **F)** Storage Modulus values at 110 s of the resins used for consecutive printing, n=3, mean ± SD., results were significant at p ≤ 0.05 for before printing vs. 8^th^ print and 1^st^ vs. 8^th^, Kruskal-Wallis; **G)** Resolution of the printed samples positive features after consecutive printing n≥3, mean ± SD., results were significant at p ≤ 0.05 for 1^st^ vs. 8^th^ print, Kruskal-Wallis; **H)** Vol3DP objects obtained from the reused resin, the numbers correspond to the print count; **I)** Schematic depicting cell seeding onto hydrogel discs; Microscopic photos of the hydrogel surface with attached EA.hy926 cells after **J)** 3 days and after **K)** 7 days of culture; **L)** Cell viability after 1, 3 and 7 days, n=8, mean ± SD., p < 0.05, one-way ANOVA with Dunn-Sidak post-hoc test; **M)** Z-stack images (i-iii) of the EA.hy926 cells attached and growing on the surface of hydrogel discs, cell nuclei are labelled in blue and F-actin in green, scale bars correspond to 50 µm.

During Vol3DP, the entire hydrogel volume is exposed to light, resulting in an anisotropic 3D dose distribution throughout the resin [45]. Unpolymerized resin can be recovered and potentially reused for new prints, but prior light exposure may cause off-target grafting (**Figure 3B**). Repeated light exposure during printing can alter the resin’s crosslinking behavior. Rheological analysis of the resin over eight consecutive printing cycles (**Figure 3C**) revealed a gradual increase in curing time. During the first print, crosslinking began 15 seconds after light exposure. The curing time increased by 1-3 s per cycle, reaching ∼27 s by the eighth print. This reduced curing efficiency is likely due to radical cleavage or partial depletion of radicals resulting from repeated light exposure of the photoinitiator [29]. Oxidative degradation of polymer networks, potentially enhanced by repeated exposure to atmospheric oxygen during resin handling, may also contribute to reduced crosslinking efficiency and mechanical performance [51]. Following the first three prints, the maximum G’ values achieved by the resins after 110 s of exposure decreased by 5-8%, but remained stable, indicating consistent performance during these cycles (**Figure 3E**). However, from the fourth printing cycle onwards, a decline in G’ was observed, reaching ∼45 %. This G’ decrease continued during the next prints, resulting in G’ differences of ∼1400 Pa between the first and the last prints. In addition to the partial depletion of the LAP [29], chemical alterations in the resin during successive printing cycles likely contributed to a decline in G’ values. However, we have excluded the potential impact of heating and cooling cycles on resin photocrosslinking abilities (**SI, Figure S2-B**).

Additionally, over the eight printing cycles, the minimum dose of light required to print an 8 mm diameter star-shaped part increased from 585 ± 17 to 825 ± 21 mJ/cm^2^ (**Figure 3D**), corresponding to the observed increase in crosslinking time with each print (**Figure 3C**). A gradual reduction in the size and precision of the parts was also noted (**Figure 3H**), making them more fragile to handle. The areas and perimeters of the printed objects decreased by ∼41% and 27%, respectively (**Figure 3G, SI Figure S2-D**). At the same time, the fineness (size of positive features) increased 2–3 times, indicating a reduction in resolution (**Figure 3F**).

The observed ∼45% reduction in storage modulus after eight consecutive prints highlights the practical limitations of reused resins. This deterioration was accompanied by increased curing times and loss in print resolution and structural integrity, particularly after the fourth cycle. Although blending recovered resin with fresh material at 1:1 or 1:2 ratios partially mitigated the loss in mechanical performance in preliminary tests, these findings underscore that direct reuse introduces variability. Resin reuse reduces material costs, preparation time, and waste, which is particularly valuable when using complex or expensive formulations. Future work should explore more robust regeneration strategies, such as photoinitiator replenishment or oxidative stabilization, to enable consistent performance in reuse scenarios.

### 2.4 Biocompatibility of the Photocrosslinked Hydrogel Constructs

The effects of hydrogel on cell survival were investigated. To determine if cells can attach to hydrogel discs made of resin, a cell suspension was introduced into a well with hydrogel discs at its bottom (**Figure 3I**). We first noted a small number of EA.hy926 cells scattered on the disc, which then began to aggregate and spread across the surface (**Figure 3J-K**). Cell viability assays demonstrated the biocompatibility of the of the photocrosslinked hydrogel constructs derived from the resin. with consistently high viability percentages (>80%) from day 1 through day 7 (**Figure 3L**), indicating sustained and improved cell proliferation over the observed period.

EA.hy926 cells were seeded onto 2D hydrogel discs (diameter 5 mm, thickness 0.6 mm) fabricated from the optimized GelMA–PEGDA resin, and viability was monitored over 7 days using the CellTiter 96® Aqueous One Solution (MTS) assay. Day 1 viability values were set to 100%, so proliferation over time resulted in values exceeding 100% (**Figure 3L**). The MTS assay was selected for its compatibility with adherent cells; potential optical interference from the hydrogel was minimized by using thin discs (≤1 mm) and removing media before absorbance reading.

Fluorescent z-stack imaging was performed on discs to observe cell morphology at different focal planes. Cells at the bottom exhibited elongated morphologies, indicative of robust integrin-mediated adhesion facilitated by gelatin-derived bioactive peptides (**Figure 3M-iii**) [47]. In contrast, upper-layer EA.hy926 cells exhibited more rounded morphologies (**Figure M-i, ii**), likely due to limited direct interactions with the substrate and possible diffusion-limited availability of adhesion factors and nutrients.

Previous reports have noted that cells adhere and proliferate better within PEG and GelMA gels compared to those composed solely of PEG [52]. The EA.hy926 endothelial-like cells adhered to the inner surfaces and remained viable over time (**Figures 3K, L).** However, full circumferential endothelialization was not assessed in this study. Confocal cross-sections or volumetric imaging would be required to confirm complete lumen coverage. In future work, we aim to employ dynamic seeding protocols and 3D imaging to ensure and validate uniform endothelial lining across the entire vascular lumen, which is essential for functional vascular tissue engineering applications. In this study, EA.hy926 cells were not encapsulated within the Vol3DP constructs. The GelMA–PEGDA formulation (5%/10%) was optimized for print resolution and mechanical stability, but not for cytocompatibility during photopolymerization. It is also important to note that EA.hy926 cells, which are frequently used in vascular research due to their stability and ease of handling, are a hybrid endothelial cell line derived from HUVECs and A549 carcinoma cells. As such, they do not fully recapitulate the gene expression profile, morphology, or functional responses of primary endothelial cells. In this study, EA.hy926 cells served as a robust model to establish proof of concept for perfusion compatibility and basic cytocompatibility with the printed constructs.

### 2.5 Prototyping Branched Vascular Structures with Vol3DP

We investigated how the resin can be utilized to prototype branched vascular structures (**Figure 4A**) to mimic hierarchical branching patterns characteristic of physiological vasculature. The resin’s low gel strength limits the fabrication of standalone vascular geometries by Vol3DP. We employed Vol3DP to pattern channels in a 3D hydrogel matrix (**Figure 4A-B**). Consequently, minimal channel wall thicknesses were maintained above 500 µm to avoid structural collapse during post-processing and handling. The perfusability of the branched channels was validated qualitatively through successful dye filling (**Figure 4C**, **SI-video-S2**) and quantitatively confirmed by perfusion tests, which ensured consistent fluid movement without observable obstructions at flow rates approximating physiological values (∼1–5 mL/min) (**SI-video-S14**). These values correspond to shear rates ranging from 1600 s^-1^ to 100 s^-1^ at which rates the viscosity of blood is nearly constant [53].

**Figure 4.**
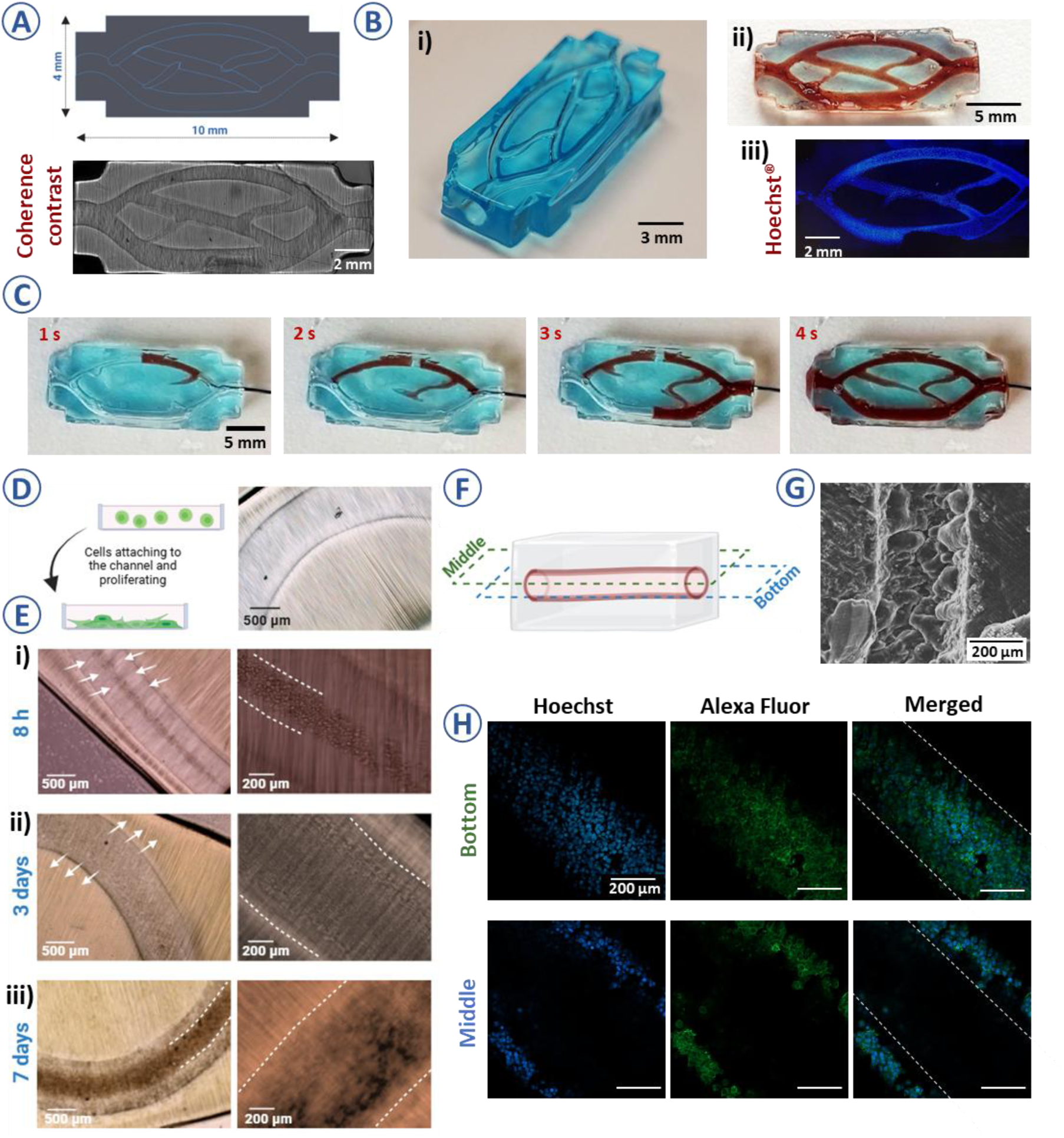
**A)** A proof of concept design of a branched vascular network (i) and coherence contrast microscopy image of the resulting, Vol3DP geometry (ii) **B)** Photograph of the embedded vascular patterns, after removing unpolymerized resin and filling the channels with (i) air (ii) red-colored PBS and (iii) fluorescent tile image of Hoechst-labelled EA.hy926 cells grown within the channels (day 7); **C)** Hand-guided perfusion experiments; **D)** Schematics of the EA.hy926 cells settling at the channel’s bottom and the image of Vol3DP channel prior to cell seeding; **E)** Vol3DP channel with seeded EA.hy926 cells (i) after 8h of incubation, (ii) after 3 days, EA.hy926 cells are more dispersed and uniformly covering the lumen, (iii) after 7 days, EA.hy926 cells start to accumulate at the bottom of the channel; **F)** Schematic of the Vol3DP vascular model showing the middle and bottom layer of the channel at which the imaging was performed; **G)** SEM image of the sample crossection revealing the channel; **H)** Images of the fluorescently labelled EA.hy926 cells (day 7) lining the Vol3DP channel, image taken in the middle of the structure reveals presence of EA.hy926 cells on the walls, shaping the lumen.

Volumetrically printed networks were later seeded with EA.hy926 cells (**Figure 4E, SI, Figure S3**). After 8 h, the injected EA.hy926 cells sedimented at the bottom of the channels (**Figure 4E-i**). To improve uniformity, rotating or flipping the constructs up to a few hours post-seeding improved initial cell distribution [11]. To aid microscopic observations after staining samples were cut in half as depicted in **Figure 4F-G**. After 3 days of culture, cell spreading was observed, with EA.hy926 cells migrating from the bottom of the channels to the sides, making the bottom layer less distinct (**Figure 4E-ii**). Similar observations were reported by Polacheck et.al. in fibronectin-treated PDMS channels [11]. After 7 days, we observed that EA.hy926 cells populations began to accumulate at the bottom of the channels, likely due to an increase in cell density and gravitational effects. This resulted in less uniform cell distribution and the formation of dense clusters, which obstructed light transmission during microscopy (**Figure 4E-iii**). Confocal z-stack imaging (**Figure 4H**) confirmed cell presence within the channels. The bottom and top of the channels were densely populated with EA.hy926 cells, while the middle lumen exhibited cell attachment primarily along the edges (**Figure 4H**). However some gaps in coverage were visible, additional confocal z-stack imaging at multiple time points and orientations is planned in future works to confirm whether these gaps represent true absence of cells or sampling artifacts. Fluorescent staining of cell nuclei (**Figure S3**) further demonstrated colonization of branched vascular geometries.

### 2.7 Perfusion Platform for 3D Printed Vascular Models

Although 3D-printed vessel models are generally more complex than widely used microfluidic technologies, the apparent advantage is that 3D fabrication technology is closer to 3-dimensional physiological conditions, which may not always be reflected in flow studies using PDMS microfluidic chips [54, 55].

Volumetric printing can create standalone or embedded vessels. However, the final geometry of the sample sets constrains the assembly of the perfusion setup. In this work we focused on addressing these geometry-related assembly constraints by (i) designing inlet and outlet solutions for different models, (ii) ensuring feasible structural support during testing and (iii) adjusting flow conditions (flow rates 0.5-15 mL /min) to achieve specific criteria such as generating the desired shear stress at the vessel walls (**Figure 5A**). To control the flow within Vol3DP structures, support platforms were designed using CAD software and fabricated using Digital Light Processing (DLP) technology (**Figure 6B-C**). An example of the setup with a peristaltic pump is shown in **Figure 5B-ii**.

**Figure 5.**
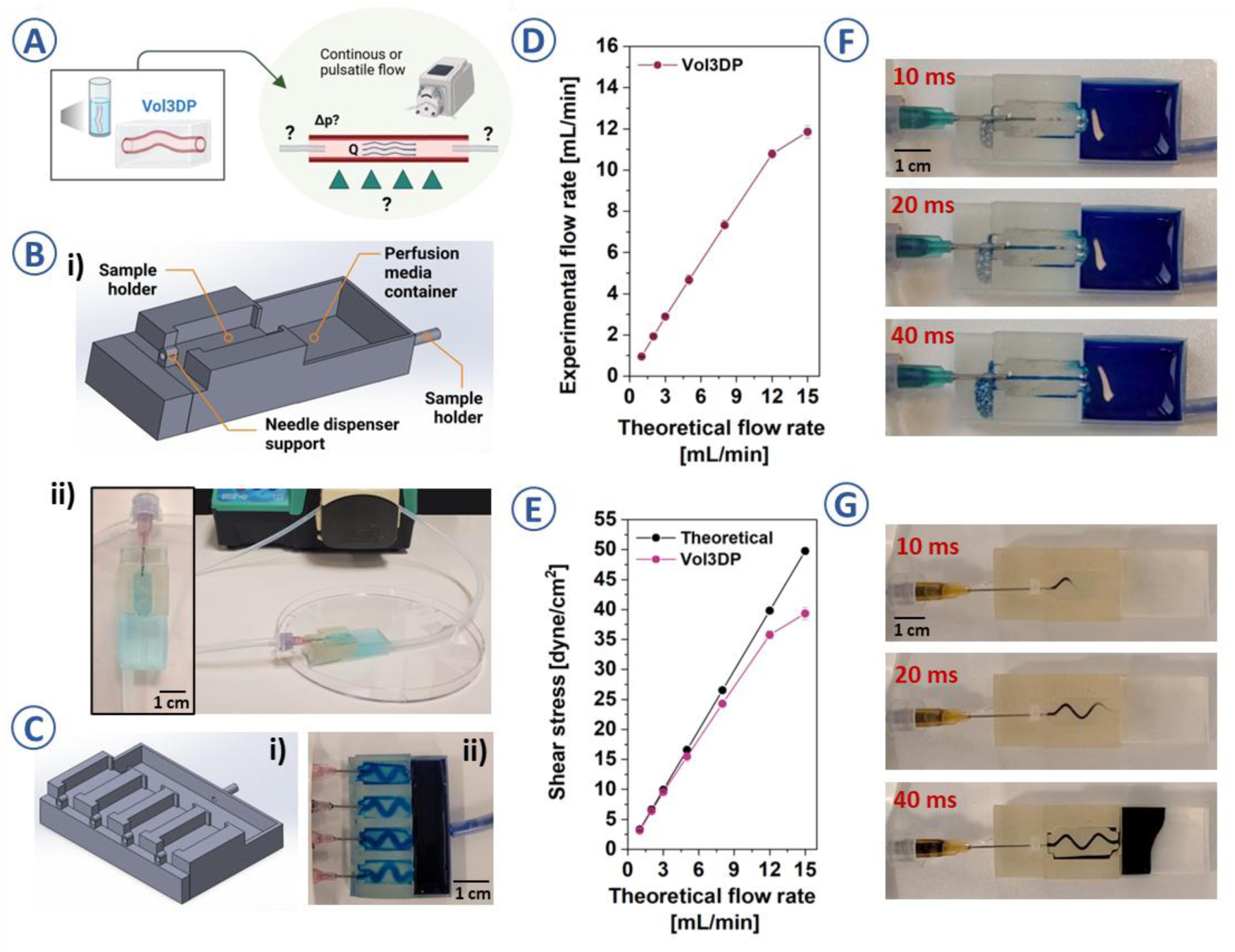
**A)** Schematic representation of the vascular geometries achievable within Vol3DP (standalone channels), crucial factors to consider in designing the model-relevant perfusion platforms; **B)** The platform schematic designed for the peristaltic pump-driven perfusion of Vol3DP channels (i). Besides, the needle holder platform is equipped with a fluid recovery container that can be directly connected to tubing, creating a closed circuit for long-term perfusion (ii) Example of the setup assembly for peristaltic pump-controlled perfusion studies for Vol3DP models; **C)** Scaled-up platform for simultaneous perfusion of up to four Vol3DP vascular models, (i) design schematic, (ii) assembled setup; **D)** Theoretical and experimental flow rate dependency for the Vol3DP models **E)** The calculated shear stress values for the employed models; Examples of the perfusion studies for Vol3DP vascular models. Vol3DP gives higher freedom of channel geometries, easily creating straight **(F)** or sinusoidal **(G)** channels;

**Figure 6.**
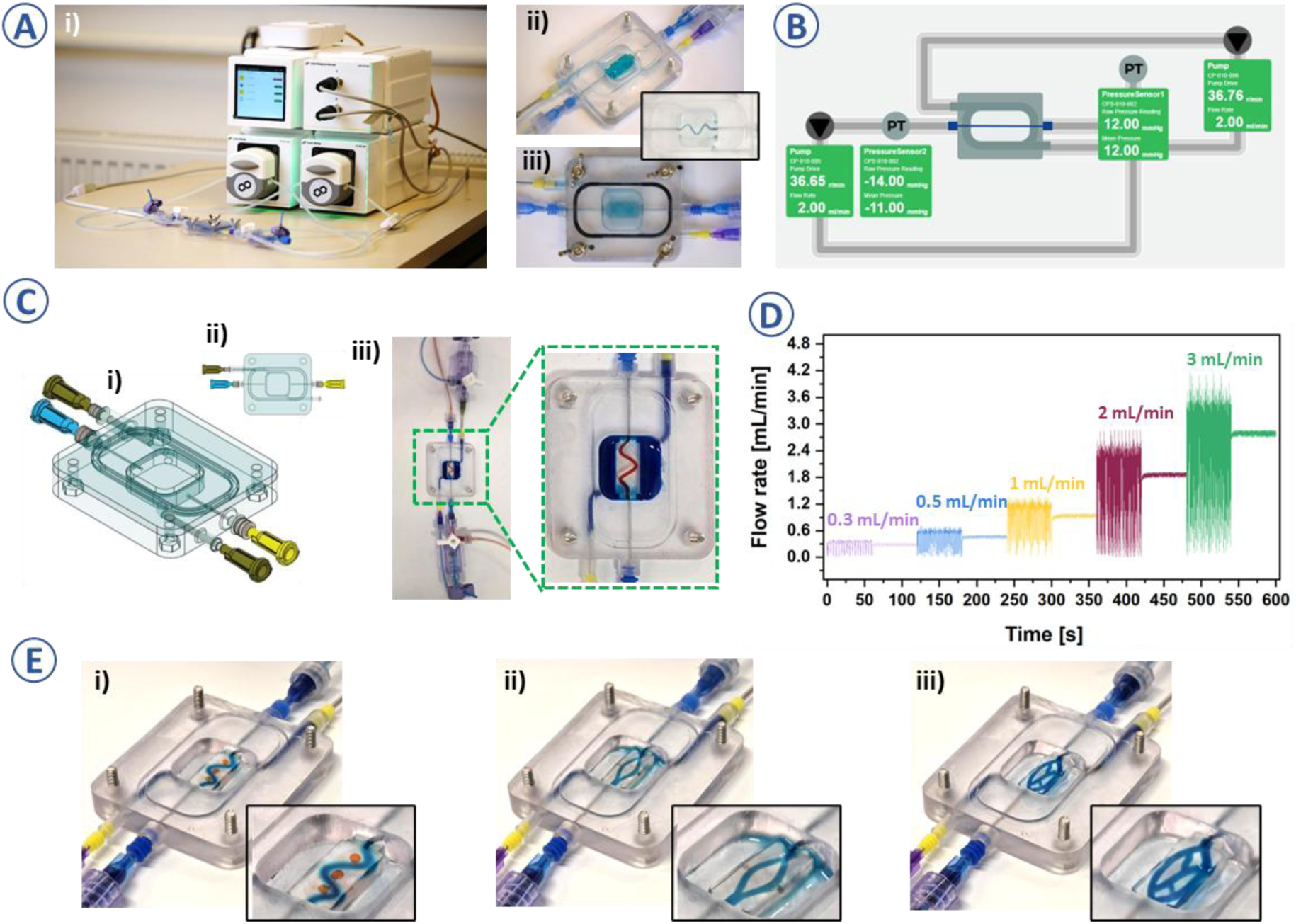
**A)** Integrated closed chamber perfusion setup for Vol3DP models testing (i), chamber assembly with Vol3DP model (ii, iii); **B)** System’s graphical interface designed for easy control of fluid flow rate and pressure monitoring; **C)** Chamber design for Vol3DP model side view (i) and top view (ii). Perfusion experiments carried out within the assembled setup (iii); **D)** Flow rate measurements for the perfusion system. The set flow rates are plotted in different colors, where the first measurement was carried out without a pulse dampener and the second one with a pulse dampener; **E)** Perfusion chamber adapted for Vol3DP sample testing with high freedom of model design e.g., sinusoidal (i), branched (ii) or hierarchical vascular-like network designs (iii).

A key requirement for hydrogels used in vessel tissue engineering is their ability to withstand hydraulic pressure and maintain continuous fluid flow [56, 57]. Recent studies have reported achieving perfusion using a laboratory rocker-shaker that leverages repeated angular shifts to induce gravity-driven flow. A key advantage of this approach is that multiple vascular platforms can be placed on the same rocker to introduce the flow [11]. However, such a gravity-driven flow method is limited in its ability to fully perfuse vascular models shorter than 3-4 mm, due to the restricted tilting angle of the rocker and the dominant capillary forces that affect fluid perfusion. Assessing successful flow remains challenging, as validation relies on indirect indicators such as changes in chamber volume or the movements of air bubbles. To address the limitation of the rocking platform, we employed a peristaltic pump to introduce controlled hemodynamic flow (**Figure 5B-ii**), minimizing the capillary effects and improving flow control.

Previous studies, including those by Bernal et al. [1], have emphasized the importance of fluid-driven stimulation for biologically relevant 3D structures. However, their approach of using a box-like chamber to expose printed constructs to bulk flow did not establish a direct connection between the Vol3DP structure and a directional flow path. Developing a perfusion platform for vascular models within this study was preceded by multiple design prototypes, which helped to identify experimental challenges. To ensure precise fluid delivery and prevent needle displacement, needle support was added. Finally, the outlet part of the platform was equipped with a fluid recovery container that can be directly connected to tubing to enable media circulation (**Figure 5B**). To increase experimental efficiency, the perfusion platforms were refined to accommodate up to four Vol3DP vascular models simultaneously (**Figure 5C**). By incorporating a tubing splitter and refining the platform design, a single peristaltic pump could perfuse four parallel channels.

We performed peristaltic pump-driven flow studies on Vol3DP platforms, testing flow rates ranging from 1 to 8 mL/min for up to three days. The system is compatible with longer-term experiments, provided that the media is replenished periodically (**SI, video S3**). For the testing purposes we considered straight-vessel models and sinusoidal channels (**Figure 5F-G**). Following **Eq. 10-11,** we adjusted flow rates to achieve physiologically relevant shear stresses of 3-50 dyne/cm^2^ within the vascular lumens [58].

For flow rates above 5 mL/min, complete elimination of leakage at the needle-model interface was not possible. However, at low flow rates (<5 mL/min), leakage had a minimal impact on flow stability (**Figure D-E).** At higher flow rates (> 12 mL/min), leakage increased, resulting in greater differences in the theoretical and experimental flow rates as measured by the collected fluid volume at a given time **(Figure 5D**). Consequently, theoretical and experimental shear stress values differed by approximately 20 % for Vol3DP at a flow rate of 15 mL/min.

Nonetheless, flow rates around 1 mL/min are sufficient to introduce dynamic conditions into the cell culture, generating shear stresses greater than 3 dyne/cm², which have been shown to promote the development of functional vascular endothelium [11]. Additionally, most perfusion culture studies operate within a flow rate of 0.1–3.0 mL/min, as excessively high flow rates have been associated with reduced cell viability [55]. Moreover, our setups can generate high shear stresses (∼40 dyne/cm^2^), proving their relevance beyond mimicking shear stress in large veins (∼1 dyne/cm^2^) and suitability for mimicking flow in small arterioles, where shear stress can reach 50 to 80 dyne/cm^2^ [58] (**Figure 5E**).

Although this study focuses on the design and implementation of perfusion platforms tailored to the unique geometries of Vol3DP constructs, we acknowledge the absence of benchmarking against commercially available perfusion platforms, such as the AIM Biotech and Kirkstall Quasi Vivo systems [59, 60]. These systems offer standardized flow environments and are optimized for simpler geometries. In contrast, our platforms enable customized perfusion with branched and tubular geometries, as well as pressure and shear stress modulation.

### 2.8 Precision Control Over The Flow Parameters and Diffusional Permeability

Subjecting a tissue scaffold to continuous perfusion necessitates the ability of the scaffold to withstand mechanical forces resulting from pumping, i.e., pressurization and fluid shear, as well as platforms that enable the control of these parameters.

Originating from our previous open platform designs (**Figure 5**), we engineered a closed-chamber adaptable platform capable of accommodating 3D-printed models to evaluate the performance of the construct-platform assemblies under flow conditions (**Figure 6A-C, SI video S4**). We applied flow rates ranging from 0.3 to 3 mL/min and included uni- and bidirectional flow conditions, which could be modulated using the graphical interface of the modular cube setup (**Figure 6B**). Due to the limited throughput of the 3D model, flow rates above 4 mL/min were not feasible, as they led to hypotensive conditions within the closed chamber.

To our best knowledge, this platform uniquely allows for simultaneous precision control of fluid flow within the vascular lumen and around its external environment (**Figure 6C, SI video S4**). Such pressure gradients in extracellular matrix (ECM) have been found to modulate sprouting angiogenesis [61] and cell migration [14].

Although flow and diffusion control are standard in hydrogel microfluidics, our system integrates these features into freeform, 3D-printed vascular constructs using Vol3DP. This enables non-planar, branched, and tunable geometries that surpass traditional chip-based platforms. The reported diffusion results are specific to the GelMA– PEGDA resin and may vary with other formulations. Additionally, the designed platform could be adapted for hypoxia studies by sealing the chamber and controlling gas exchange in future work.

Previous studies have indicated that morphological changes in blood vessels occur at shear stress levels of approximately 5 dyne/cm², corresponding to flow rates of 1-2 mL/min in our systems, which contributes to the formation of a functional vascular barrier with reduced leakage. This provides a foundation for further iterations using cell-embedded constructs, where hydrogel-embedded EA.hy926 cells could be used to quantify vascular permeability or investigate cytoskeletal changes [11] under different flow conditions.

Unlike traditional shear chambers, our assembled setups employ pulsatile flow, inducing both pulsatile strain and the expansion/contraction of the surrounding hydrogel, making the model sensitive to pressure in addition to shear stress, mimicking the systolic and diastolic pressure dynamics characteristic of physiological blood flow [11].

However, peristaltic pumps can be adapted by incorporating pulsation dampeners to study laminar flow conditions when necessary [62]. Pulsatile flow was achieved by operating the peristaltic pump without the inline pulse dampener, transmitting its inherent cyclic pressure waveform directly to the construct. Previous studies have shown that continuous laminar flows in perfusion culture promote uniform endothelial cell alignment and junction formation. At the same time, pulsatile flow tends to result in more randomized cell organization [62, 63]. In our setup, the Integration of a pressure dampener within the inlet tubing effectively smoothened the flow profile and functioned reliably across various flow rates (0.3-3 mL/min) (**Figure 6D**). Therefore, the developed platform is capable of delivering both pulsatile and continuous flow, supporting its versatility and potential in blood flow mimicry and perfusion-based cell culture, respectively.

The geometries presented in this study are relatively simple (**Figure 6E, SI videos S7 and S8**) and serve as controlled models to validate perfusability, fabrication fidelity, and cell compatibility. We acknowledge that such designs could also be achieved with other techniques, such as sacrificial molding or lithographic patterning. However, the intrinsic ability of Vol3DP to fabricate complex, non-layered, freeform geometries within seconds remains beneficial for rapid prototyping and iterative design.

As a preliminary proof-of-concept, we employed methylene blue dye diffusion to assess the relative permeability of our vascular models [11] (**Figures 7A-B, SI Figure S4, SI Videos S4 and S5**). We observed the dye spread through the channel walls, with approximately 30% diffusion occurring within 30 minutes (Figure 7B-ii), ultimately resulting in a dye-covered area of approximately 1 cm² (**Figure 7B-iii**). Although this approach offers visual and semi-quantitative insights into solute transport across the hydrogel walls, it does not capture size-selective permeability or precise barrier integrity. This can be more accurately evaluated using molecular tracers, such as FITC-dextran (particularly the 70 kDa variant), which is widely considered the standard for assessing endothelial permeability and paracellular transport in vascularized constructs. However, methylene blue was selected here for its strong visible absorbance, low cost, and rapid imaging capability for proof-of-concept testing; future work will use FITC-dextran for quantitative assessment.

**Figure 7.**
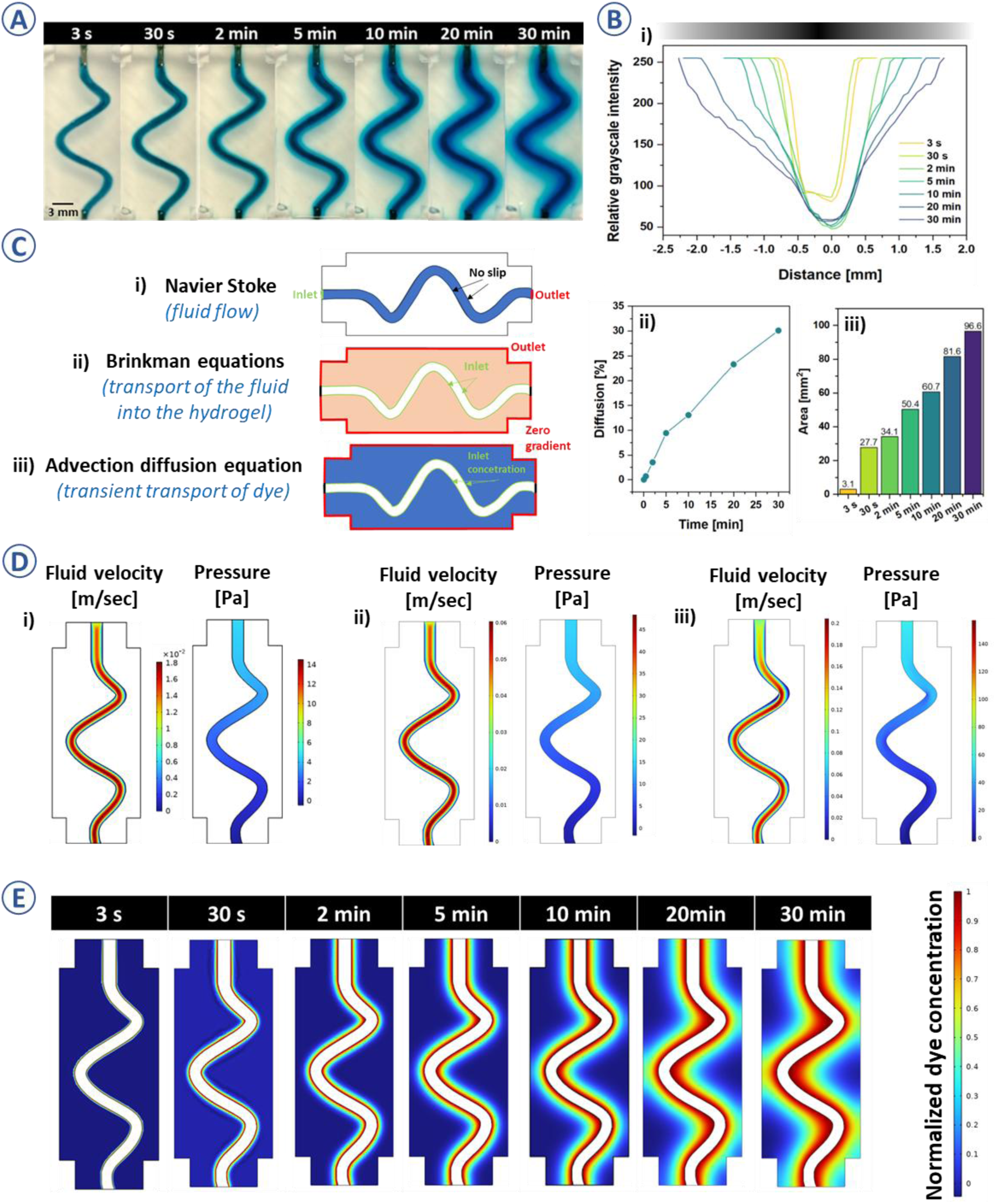
**A)** Dye permeation images taken at different perfusion timepoints; **B)** Dye permeation distance plotted against relative grayscale intensity of the taken image allowing to distinguish higher intensity within the channel and lowering intensities of permeation areas, for the flow rate 1 mL/min (i), Quantified % of diffusion into the channel walls (calculated as percentage intensity) (ii); Increase in the dye coverage area of the sample (iii); **C)** Geometry and boundary conditions for the different solved problems using computational simulation: (i) In fluid flow analysis, the perfusion comes from the left, with a constant velocity (inlet) to the right (outlet), (ii) To simulate fluid transport inside the hydrogel, the Brinkman’s equations are solved, (iii) To simulate dye transport, advection diffusion equations are solved, with the vessel walls as the inlets; **D)** Fluid velocity magnitude (m/s) and pressure (Pa) inside the vessels for the three different simulated fluid flows (i) 0.3 mL/min, (ii) 1 mL/min, (iii) 3 mL/min; **E)** Relative concentration of dye in the porous medium compared to that in the artificial blood vessel at different time points for fluid flows of (i) 1 mL/min, dimensions of the simulated gel and embedded vessel were taken from the 3D-printed geometry.

### 2.9 Computational Simulations of Flow and Dye Transport

To complement our experimental assessment of perfusion and permeability, we performed computational simulations using COMSOL Multiphysics (**Figure 7C-E**). Transient two-dimensional (2D) simulations were conducted to investigate fluid flow and dye transport dynamics in the printed vascular constructs. The model separately solved Navier-Stokes equations for laminar flow in the vessel and the Brinkman equations for fluid penetration into the porous hydrogel, assuming negligible hydrogel perfusion compared to intraluminal flow. Following the resolution of the flow field, dye transport into the hydrogel matrix was simulated using the transient advection-diffusion equation. Water was modeled at 25 °C with a density of 997 kg/m³ and dynamic viscosity of 1.002 mPa·s. The hydrogel was treated as a porous medium with a characteristic pore size of 100 nm, corresponding to a permeability of ∼1.0 × 10⁻¹^4^ m² based on the relation by Cuccia et al. [64]. The porosity (0.6) and dye diffusivity (6.74 ⋅ 10^−10^*m*^2^/*s*) were set to standard values for polymer hydrogels and methylene blue, respectively.

Boundary conditions (**Figure 7C**) included constant inlet velocity for vessel perfusion, zero-gradient outlet conditions, and pressure-matched walls between the vessel and hydrogel. For dye transport, the vessel wall served as a concentration source, with inlet values matching those in the luminal fluid.

Simulations were conducted at three flow rates of 0.3, 1.0, and 3.0 mL/min, corresponding to Reynolds numbers ranging from 8 to 80. These simulations confirmed the laminar regime while revealing an increasing asymmetry in the velocity distribution at higher flow rates due to curvature-induced effects (**Figure 7D, E**). Pressure gradients scaled proportionally with flow rate. Despite this, dye permeation into the hydrogel showed minimal dependence on the flow rate (**Figure 7E, SI, Figure S5, videos S9-11**), consistent with the experimental data (**Figure 7A**). This suggests that dye transport was primarily driven by diffusion rather than convection, with only modest enhancement from advective effects at higher flow rates.

Overall, the simulations confirmed experimental observations and validated the modeling approach. These results underscore the dominance of diffusive transport through the hydrogel matrix and highlight the ability of our system to simulate physiologically relevant mass transfer phenomena.

### 2.10 *In Vivo* Validation of Flow-Aided Dye Distribution Using a Larvae Model

To complement our *in vitro* perfusion studies and highlight the physiological relevance of dynamic flow, we conducted a pilot investigation using *Galleria mellonella* larvae to contrast biologically active and passive dye transport. We frame larvae experiments as an exploratory, low-cost *in vivo* analogue to illustrate the role of perfusion-driven transport, not as a direct vascular mimic. This experiment served as a simple *in vivo* analogue to assess active versus passive transport phenomena, complementing the *in vitro* diffusion data presented earlier. Methylene blue was injected into the proximal segment of live and dead larvae (**Figure 8A**), and dye migration was monitored at 1, 5, and 10 minutes post-injection by quantifying absorbance at 665 nm across three anatomical segments: proximal, middle, and distal.

**Figure 8.**
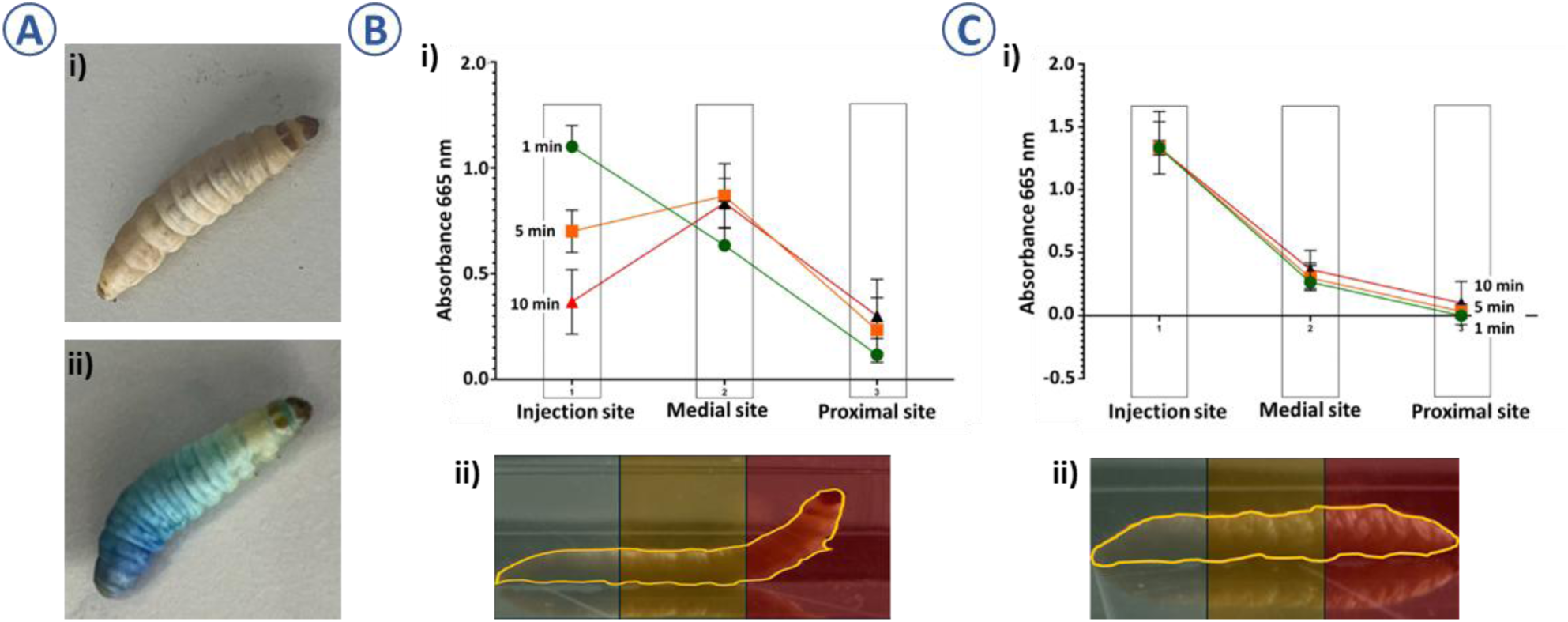
**A)** Non-injected larva (i) and larva injected with methylene-blue dye (ii); **B)** Segmental and temporal distribution of methylene blue (i) in G. mellonella larvae - alive (ii), **C)** Segmental and temporal distribution of methylene blue (i) in G. mellonella larvae - killed by liquid nitrogen (ii); Larvae were injected with 10 µL of 10 mg/mL methylene blue into the proximal segment. Absorbance at 665 nm was measured in the injection, middle, and distant body segments (depicted in the lower panel of B and C panels) after 1, 5, and 10 minutes. In live larvae, the dye propagated over time with a peak concentration shifting toward the middle segment. In dead larvae, diffusion was limited and remained mainly in the proximal region.

As shown in **Figure 8B**, live larvae exhibited dynamic redistribution of the dye, with a clear shift of peak absorbance toward the middle segment over time. This suggests an active transport mechanism likely facilitated by hemolymph circulation or muscle-driven peristalsis. Dye concentration did not equalize uniformly across all segments, implying non-linear flow characteristics and possibly retrograde transport. In contrast, dead larvae (**Figure 8C**), euthanized via rapid freezing and lacking circulatory function, retained the dye primarily in the proximal segment. Only minimal, time-dependent passive diffusion was observed, and the dye rarely reached the distal portion, even at the 10-minute mark. These findings underscore the critical role of active biological functions in compound distribution and reinforce the importance of incorporating perfusion in engineered vascular platforms. Similar to the active redistribution observed in living larvae, our perfused hydrogel constructs enable controlled and directional solute transport, providing a more physiologically relevant model compared to static systems. This comparative insight further validates the need for perfusion-capable scaffolds in simulating vascular environments for drug delivery or permeability studies. Consequently, the distinction between live and passive models should be considered when evaluating compound dispersion or pharmacokinetics in both invertebrate models and engineered tissues.

## 3. Conclusions and Limitations

This study demonstrates the rapid fabrication (<1 minute) of perfusable vascular structures using volumetric 3D printing (Vol3DP) and a reusable GelMA–PEGDA resin. Vol3DP stands out as a promising approach for fabricating vascular-like geometries in seconds. The approach enables high-resolution, embedded hydrogel architectures with tunable dimensions and flow paths. Custom-designed perfusion platforms were developed to provide precise control over flow rate, shear stress, and pressure, key parameters for simulating physiological conditions. Although the system supports both pulsatile and laminar flow and is compatible with cell culture, the current study focused on platform validation and material performance. Experimental dye permeation data were assessed against computational simulations, showing a negligible impact of fluid flow rate on permeation. To complement *in vitro* perfusion studies, we conducted a pilot investigation in Galleria mellonella larvae as a simple *in vivo* model to assess active versus passive dye transport. Functional biological responses to flow, such as cytoskeletal alignment, barrier integrity, or gene expression, were not assessed. In the present work, only endothelial-like cells were incorporated, which we acknowledge as a limitation. Future work will address these aspects using primary endothelial cells to further establish the physiological relevance and translational potential of the platform in vascular tissue engineering and drug screening applications.

## 4. Experimental section

### 4.1 Materials and Methods

Gelatin type A (acid-cured, porcine skin; ∼300 g Bloom), phosphate-buffered saline (PBS), Methacrylic anhydride (MAA), polyethylene glycol diacrylate (PEGDA; average Mn 700), lithium phenyl-2,4,6-trimethylbenzoylphosphinate (LAP) with purity ≥ 95%, were purchased from Sigma-Aldrich (St. Louis, MO, USA). Dialysis membrane (Mw cut-off (MWCO): 14 kDa) was obtained from Membra-Cel™, (United States). Dyes: Congo Red and Methylene Blue were purchased from TCI (TCI EUROPE N.V., Zwijndrecht, Belgium), and black particles (Noir Irgalene BGL) were obtained from Ciba-Geigy AG (Switzerland). Methacrylated gelatin was synthesized following our previous protocols [57, 65].

### 4.2 Resin formulation for Vol3DP

Different resin formulations were prepared by varying the concentrations of GelMA, PEGDA, and the concentration of photoinitiator LAP (0.3 to 1 mg/mL) [40, 66]. The quantities of polymers and photoinitiators are specified in (**SI, Table S1**). For instance, to prepare resin (GelMA 5) % (w/v), PEGDA (10 % (v/v), LAP 0.3 mg/mL). Stock solutions were prepared as follows: GelMA was dissolved in PBS at a concentration of 7.5% (w/v), and PEGDA was diluted in PBS to a concentration of 30% (w/v). All stock solutions were heated to 37 °C until completely dissolved. To prepare resin for Vol3DP, stock solutions were combined in a 2:1 ratio of GelMA: PEGDA, and lithium phenyl-2,4,6-trimethylbenzoylphosphinate was added to reach a final concentration of 0.3 mg/mL. Finally, the resin was evenly mixed, and if necessary, bubbles were removed in an ultrasonic bath [67].

### 4.3 Volumetric printing – Vol3DP

Resins were dispensed into cylindrical borosilicate (BK7) glass vials (Ø 15 mm) [68]. Vials were loaded into a commercial volumetric 3D printer (Tomolite, Readily3D, Switzerland; multiwavelength). The samples were thermally gelated at 4°C prior to printing. Models for printing were designed in SolidWorks (St. Waltham, United States) and generated.STL files were loaded into the printer software (Apparite, Readily3D, Switzerland). Prior to printing, the resin’s refractive index was measured using a SmartRef (LAB Meister, Anton Paar GmbH, Graz, Austria) and input into the software. Printing was performed with a 405 nm light source. After printing, the vials were heated to 37°C, and uncured resin was collected. The printed constructs were then washed gently with 37°C PBS [1, 28].

### 4.4 Characterization

#### Resolution test

To evaluate the resolution capabilities of the volumetric printing technique, we have designed two models. The first consisted of a star-shaped object with a diagonal 3 mm and a thickness of 1 mm, designed to assess the minimum feature size - fineness (a positive characteristic) achievable with the printing process (**Figure 2G**). The second model was a rectangular block (8 mm width, 4 mm thickness, 15 mm height) incorporating internal channels with diameters decreasing from 1.5 mm to 0.1 mm to determine the minimum printable internal diameter (a negative characteristic) (**Figure 2F**). The models were printed and measured to compare the model and print dimensions.

#### Rheological properties

Using an Anton Paar MCR 302 rheometer (Anton Paar, Ghent, Belgium), the crosslinking kinetics of the resins were assessed. Time sweep experiments were performed at a frequency of 10 Hz, with 1.0% constant strain at 25°C (n = 3, independent measurements); 15 seconds after the start of the measurement, the light source (Dymax, QX4, VisiCure – 405 nm, intensity: ∼14.9 W/cm², Mavom, Kontich, Belgium) was activated for the remaining 105 s. The plate-plate measuring system was used, featuring a 25 mm diameter upper plate and a 100 µm gap size [69, 70]. Samples were *in situ* formed on the rheometer plate with a 1 mm gap at 25 °C before starting measurements.

Temperature sweep: Experiments were performed at a frequency of 1.0 Hz, with a strain rate ranging from 0.1% to 100% strain, starting at 15°C and finishing at 40 °C. The points were recorded every 30 s.

#### Material opacity

To determine the opacity of the resins, absorbance measurements were carried out using a UV multimode plate reader (Spark, TECAN, Trading AG, Switzerland). To obtain the percentage opacity of a material from its absorbance, **Eq. 2-4** was used [71]:

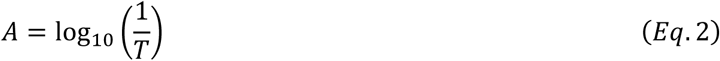

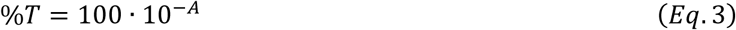

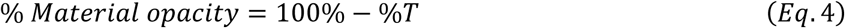

Where A is absorbance and T is the transmission.

#### Digital Light processing for rapid prototyping of sample holders and perfusion platforms

Perfusion platforms for securing objects during perfusion were designed in CAD software (SolidWorks, St. Waltham, United States). Lithographic printing was performed using the Asiga MAX ASIGA 27X DLP printer (Erfurt, Germany), equipped with a 405 nm wavelength UV LED light projector. Parts were printed using ABS-Like Resin+ (Shenzhen Anycubic Technology Co., Ltd.), and Asiga Composer software was used to position the CAD models on the build platform and set parameters (layer thickness of 100 µm, layer exposure time of 10 seconds, and light intensity of 15 mW/cm²). After 3D printing, the parts were washed with isopropanol to remove all resin residues [72].

#### Microscopic evaluations and sample imaging

Images of the resins and 3D-printed samples were taken using a Fujifilm digital camera (Fujifilm X-H2S). Methylene blue was used to stain the printed constructs, allowing for better visualization of the otherwise transparent hydrogels.

3D printed samples were examined under an inverted microscope (Zeiss Axio Observer, Germany, 5x and 20x objectives) to measure the diameter of printed vessels and assess the finesse of printed structures. We used the coherence contrast mode of the microscope. A minimum of three measurements was taken per sample to calculate average values. Measurements were performed using ZEISS Zen software [28].

Brightfield images during cell culture were taken using a Cell Imager (ZOE, Bio-Rad, Hercules, CA, United States). Fluorescently labeled samples were observed under a ZEISS LSM900 confocal inverted microscope using Airyscan mode (Zeiss, Germany). Micrographs were taken at randomly selected positions to qualitatively assess morphology and distribution of cells [73].

Scanning electron microscope (SEM) images were taken to investigate cell morphology and adhesion for different hydrogel samples (FEI Quanta 200 FEG). For this, cell-seeded scaffolds were fixed with a 2.5% glutaraldehyde solution in cacodylate buffer, and subsequently dehydrated with a graded ethanol series (50%, 75%, 90%, and 98%). Samples were air-dried for 3 days, then cross-sectioned and coated with platinum before being observed under SEM [57, 74].

#### Cell culture and viability assay

Modified endothelial cell line - EA.hy926 was maintained with Dulbecco’s minimal essential medium (high glucose), supplemented with 10 % FBS, 100 µg/mL Penicillin, 100 U/mL Streptomycin (Thermo-Fisher Scientific, Massachusetts, USA), in a humidified CO_2_ chamber (37°C, 5 % CO2).

To evaluate the hydrogel’s biocompatibility and ability to serve as a support for cell growth and proliferation, EA.hy926 cells were cultured on the hydrogel discs (seeding density: 50,000 cells/cm²) for 1, 3, and 7 days. Cell viability was assessed using the MTS assay (CellTiter 96® Aqueous One Solution Cell Proliferation Assay, Promega, Madison, WI, United States). The samples’ absorbance was read at 492 nm using a microplate reader (Epoch, BioTek, United States). EA.hy926 cells viability was calculated over time (1, 3, and 7 days) based on the absorbance values of control wells containing EA.hy926 cells grown in the absence of hydrogel (**Eq. 5**).

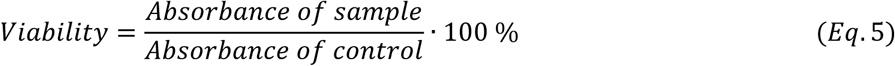

Metabolic activity was measured with a resazurin assay (resazurin sodium salt, Sigma Aldrich, St. Louis, United States) and cell proliferation was evaluated after 1 and 7 d (n = 8) [27].

#### Channel seeding

After (8-24 h) preincubation, the medium vascular models were removed from the incubator. The medium was aspirated and a 200-μL pipette was used to gently clean the trapped medium out of the channels, filling them with air instead [11]. EA.hy926 cells were dispersed at a seeding density of 2 million/mL, and the channels were gently filled with 20-25 μL of the cell suspension, ensuring that the air was fully pushed out of the channels. Once the seeding was complete the vascular models were returned to the incubator. After 24-30 h of incubation, non-adherent EA.hy926 cells were removed by gently perfusing the channels with fresh medium. This also helped to remove cell debris from the vascular models [11]. Samples were cultured for one week, with the media refreshed daily.

#### Cell staining

Cytoskeleton staining (F-actin) was performed with 10 μg/mL Hoechst and 2 μM Alexa Fluor 488 – Phalloidin. To evaluate cellular morphology, EA.hy926 cells were fixed with 4% Paraformaldehyde (PFA), for 30 min, followed by staining with 10 µg/mL Hoechst and 2 µM Alexa Fluor 488 – Phalloidin (ab176753) - Abcam, Cambridge, UK), which was performed in DPBS, 1% BSA at room temperature °C for 90 min. Samples were observed under a ZEISS LSM 900 confocal inverted microscope [57, 65].

#### Perfusion experiments

For perfusion studies involving a peristaltic pump, Vol3DP channeled structures were positioned on the DLP-printed supporting platforms (**Figure 6C-E**). The needle tip was carefully inserted and aligned with the lumen of the hydrogel. Silicone tubing (ID = 2 mm, OD = 6 mm) was then connected to the DLP-printed outlet solution and the needle tip at the inlet. Tubing was then inserted into a peristaltic pump (Watson-Marlow 323, Watson-Marlow Limited, Cornwall, United Kingdom), enabling flow control by adjusting the RPM (1-300 RPM). Using **Eq. 9** RPM were set to evoke the desired flow rate. Parts were secured in a deep-dish 100-mm Petri dish. Perfusion with PBS either pristine or supplemented with dyes; Congo Red (0.5 mg/mL), Methylene Blue (0.1 mg/mL), or black particles (Noir Irgalene BGL, 0.5 mg/mL) was performed to ease the observation of fluid flow within hydrogel structures. For the perfusion scale-up experiments, perfusion chambers Vol3DP were adapted to accommodate four samples simultaneously. To divide the fluid flow a channel splitter (RS Components, Corby, United Kingdom) was used. The tubing splitter enables one pump to perfuse four identical channels in parallel, ensuring consistent flow conditions across multiple replicates for comparative testing. A closed chamber perfusion setup was further developed by A4BEE Sp. z o.o. (Wrocław, Poland) to facilitate more accurate testing. The system was integrated with modular cubes to control flow rate and enable pressure readout. A controlled-flow perfusion system was alternatively equipped with fluid pulse dampers (SE-D1606-3NB-PC-316, Darwin Microfluidics, Paris, France) to enable the laminar flow of the medium despite using a peristaltic pump. To validate the flow rate of fluid through the construct, the outlet tubing was connected to a fluid flow sensor evaluation kit (SLF3S-1300F, Sensirion, Stäfa, Switzerland). Tubing adapters were used as needed to ensure a seamless connection of the entire setup.

##### Flow experiments using a peristaltic pump

Flow Rate was calculated using (**Eq. 10**) or directly set on the peristaltic pump. To convert the revolution per minute (RPM) into mL/min and vice versa, **Eq. 9** was used

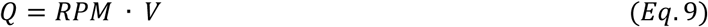

Where 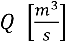 Is the flow rate, *RPM*, is revolutions per minute, and *V* [m^3^] is the volume of the fluid recovered in one minute [75].

To determine the flow rate and shear stress, we assumed viscous-dominated flow and PBS as a Newtonian fluid. We further refer to the principle of conservation of momentum for a fluid in a cylinder. Thus, we formulate the following relationship between the volumetric flow rate and the wall shear stress [11]. The transformed formula **(Eq. 11**) was used to estimate the wall shear stress.

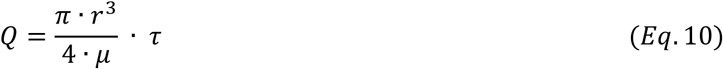

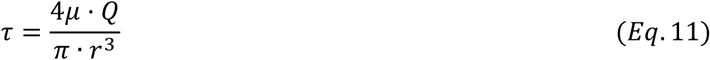

where 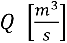 is the volumetric flow rate, r [m] is the vessel radius, μ is the fluid viscosity (µ_PBS_= 0.0089 Pa·s), and τ is the shear stress [Pa] at the vessel wall.

##### Permeability quantification

Permeability was assessed by recording the perfusion of 10 mg/mL methylene blue dye at a flow rate of 1 mL/min over 30 min through the printed channels. Images were taken at different time points (0-30 min) using a Fujifilm camera (Fujifilm X-H2S). The dye diffusion images were converted into 8-bit, grayscale images (**SI, Figure S4**) with pixel values (0-255) and plotted as a function of dye diffusion distance, over a total length of 4 mm. The percentage diffusion of methylene blue was quantified based on the average fluorescence intensities within the lumen and the channel walls **(Eq. 12)** [76].

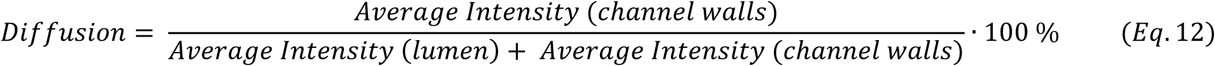

#### Numerical Simulations

Computational simulations were performed in COMSOL Multiphysics (COMSOL Inc., Burlington, MA, USA) using a 2D transient model. Separate physics interfaces were used for: (i) laminar flow in the vessel (Navier-Stokes equations), (ii) porous flow in the hydrogel (Brinkman equations), and (iii) dye transport (advection-diffusion). Water was assumed to be the fluid medium with standard physical properties at 25 °C. A hydrogel permeability corresponding to a pore size of 100 nm (∼10⁻^14^ m²) was used, as described by Cuccia et al. [64]. Boundary conditions were assigned as follows: a constant velocity at the inlet, pressure at the outlet, and no-slip at the vessel walls; pressure values at the hydrogel boundaries were inherited from the vessel solution. Dye diffusion simulations assumed inlet concentration matching the dye level inside the lumen. Simulations were carried out for three perfusion rates: 0.3, 1.0, and 3.0 mL/min. Mesh sensitivity analysis ensured numerical stability and spatial convergence. Results were visualized as flow velocity, pressure contours, and normalized dye concentrations, using geometry dimensions derived from the 3D-printed constructs.

#### Temporal and Segmental Distribution of Methylene Blue in Live and Dead Larvae Model

Final instar larvae of *Galleria mellonella* (Platform for Unique Models Application, Wroclaw Medical University, Poland) weighing 200 ± 20 mg were used. *Galleria mellonella* experiments did not require institutional ethics approval as they are invertebrates. Only healthy, cream-colored individuals with no visible signs of melanization or injury were selected. Larvae were randomly assigned to two experimental groups: live larvae and dead larvae (n=9, each). The latter were euthanized by immersion in liquid nitrogen (-196 °C) for 10 seconds and allowed to thaw at room temperature before use. A 1% (w/v) aqueous solution of methylene blue (Biomus, Lublin, Poland) was injected into the proximal section of each larva using a 10 µL Hamilton microsyringe (model 701 RN). To assess the spatial propagation of the dye over time, each larva was divided into three equal-length anatomical segments: proximal (the site of injection), middle, and distal (the farthest from the injection site). Larvae were sampled at 1, 5, and 10 minutes after injection (n = 3 per time point per group). In the live group, individuals were euthanized by immersion in liquid nitrogen at the specified time point to preserve the *in vivo* dye distribution. Larvae in the dead group were injected with dye after thawing, allowing passive diffusion to occur without active circulation or metabolism.

Each larva was manually sectioned with a sterile razor blade into its three predefined segments. Each segment was transferred into 5 mL of ultrapure water and homogenized using a bead mill homogenizer (BeadBug™ 6, Benchmark Scientific, USA) for 2 minutes at 4000 rpm. The resulting homogenates were centrifuged at 12,500 × g for 10 minutes at room temperature. A 100 µL aliquot of the supernatant from each segment was transferred to a 96-well flat-bottom microplate (Sarstedt, Germany), and absorbance was measured at 665 nm using a microplate reader (Multi-scan GO, ThermoFisher Scientific). Control measurements included a blank (ultrapure water only) and a biological control consisting of homogenized larval tissue without methylene blue. Absorbance values were corrected for background by subtracting the mean values of both controls. The extent of dye migration was evaluated by comparing absorbance values across the proximal, medial, and distal segments at each time point. Results were reported as mean ± standard deviation (SD). No statistical analysis was performed due to the pilot nature of the experiment and limited sample size.

### 4.5 Statistical analysis

All data are reported as mean ± standard deviation (SD), with the number of independent samples (n) indicated in the figure captions. All statistical analyses were performed using Origin Pro 2021.

Given the nature of the measurements performed to determine material opacity, data normality was confirmed using the Shapiro-Wilk test, followed by a two-way ANOVA with the Tukey post-hoc test. Similarly, for the comparison of shear stress at different flow rates, data normality was confirmed using the Shapiro-Wilk test, followed by one-way ANOVA with the Tukey post-hoc test.

Given the small sample sizes, non-parametric statistical methods were used to ensure the robustness of the analysis without reliance on normality assumptions. The Kruskal-Wallis test was employed for comparisons between groups in printing resolution tests (positive and negative feature size), followed by Dunn-Sidak’s test for post-hoc pairwise comparisons.

The Kruskal-Wallis test was employed for analysis involving minimum printing dose, and G’, feature size, area, and perimeter in consecutive printing experiments. In-depth pairwise comparisons were not performed due to limited statistical power.

For cell viability results, the time point was treated as an independent factor. Data normality was confirmed with the Shapiro-Wilk test, followed by one-way ANOVA with the Tukey post-hoc test.

## Author Contributions

Conceptualization J.S.S., Data curation J.S.S., I.B.L., A.T., A.J, K.P., Formal Analysis J.S.S., A.J., M.C., M.G., Investigation J.S.S., A.J., M.C., M.G., Methodology J.S.S., A.S, A.J., M.C., M.G., Project administration A.S, Resources A.S, Supervision A.S, Validation A.S, Visualization J.S.S., A.T., A.J., M.C., M.G., Writing – original draft J.S.S., Writing – review & editing J.S.S., A.S, A.J., M.C., M.G.

All authors have read and agreed to the published version of the manuscript.

## Institutional Review Board Statement

Not applicable.

## Informed Consent Statement

Not applicable.

## Acknowledgements

The graphical abstract and figure panels 1A, 1C, 1D, 3B, 3I, 4D, 4F, 5A were created with the help of BioRender.com.

J.S.S. gratefully acknowledges the support of a grant from the FNRS (J.S.S. FNRS-Aspirant, Grant No. FC 46599).

We thank the team of engineers from A4BEE Sp.z.o.o. for their technical support in the preparation of the microfluidic modules, and in particular, we thank Kinga Surmacz and Paweł Godawa for their valuable discussions.

A remotely controlled perfusion chamber with a pressure sensor and two peristaltic pump units was designed and assembled by A4BEE Sp. z o.o. (Wrocław, Poland).

## Appendix - Supplementary data

S1.1 - Experimental section

Table S1 - Formulation of biomaterial resins

S1.2 – Results

Figure S1 - Volumetric printability evaluation for various formulated resins
Figure S2 – Volumetric printing (Vol3DP) - Material and printing concerns
Figure S3 - EA.hy926 cells seeded within a multichannel Vol3DP hydrogel
Figure S4 – Grayscale and binary images used to quantify hydrogel permeability
Figure S5 – CFD simulated permeation patterns within the hydrogel

S2 – Manual perfusion of the branched vascular model

S3 - Peristaltic pump-driven perfusion studies of the Vol3DP model

S4 – Permeation test for Vol3DP hydrogel

S5 - Perfusion and permeation of complex Vol3DP model

S6 – Intraluminal and extraluminal flow for Vol3DP model

S7 - Bifurcated channel perfusion

S8 – Perfusion of a branched vascular network

S9 - Permeation simulation for the flow rate 0.3mL/min

S10 - Permeation simulation for the flow rate 0.3mL/min

S11 - Permeation simulation for the flow rate 0.3mL/min

## References

1. Bernal, P.N., et al., Volumetric Bioprinting of Organoids and Optically Tuned Hydrogels to Build Liver-Like Metabolic Biofactories. Advanced Materials, 2022. 34(15): p. 2110054.

2. Simińska-Stanny, J., et al., Hyaluronic Acid Role in Biomaterials Prevascularization. Advanced Healthcare Materials, 2024. 13(30): p. 2402045.

3. Iqbal, M.Z., et al., Breathing new life into tissue engineering: exploring cutting-edge vascularization strategies for skin substitutes. Angiogenesis, 2024. 27(4): p. 587–621.

4. Tucker, W.D., Y. Arora, and K. Mahajan, Anatomy, Blood Vessels, in StatPearls [Internet]. StatPearls Publishing: Treasure Island (FL).

5. Duivenvoorde, L.P.M., et al., Comparison of gene expression and biotransformation activity of HepaRG cells under static and dynamic culture conditions. Scientific Reports, 2021. 11(1): p. 10327.

6. Fukushi, M., et al., Formation of pressurizable hydrogel-based vascular tissue models by selective gelation in composite PDMS channels. RSC Advances, 2019. 9(16): p. 9136–9144.

7. Wang, Y., et al., Advances in hydrogel-based vascularized tissues for tissue repair and drug screening. Bioact Mater, 2022. 9: p. 198–220.

8. Zhang, Y.S., et al., Bioprinting 3D microfibrous scaffolds for engineering endothelialized myocardium and heart-on-a-chip. Biomaterials, 2016. 110: p. 45–59.

9. Song, K.H., et al., Complex 3D-Printed Microchannels within Cell-Degradable Hydrogels. Advanced Functional Materials, 2018. 28(31): p. 1801331.

10. Rapp, J., et al., 2D and 3D in vitro angiogenesis assays highlight different aspects of angiogenesis. Biochimica et Biophysica Acta (BBA) - Molecular Basis of Disease, 2024. 1870(3): p. 167028.

11. Polacheck, W.J., et al., Microfabricated blood vessels for modeling the vascular transport barrier. Nature Protocols, 2019. 14(5): p. 1425–1454.

12. Osaki, T., V. Sivathanu, and R.D. Kamm, Crosstalk between developing vasculature and optogenetically engineered skeletal muscle improves muscle contraction and angiogenesis. Biomaterials, 2018. 156: p. 65–76.

13. Linville, R.M., et al., Physical and Chemical Signals That Promote Vascularization of Capillary-Scale Channels. Cellular and Molecular Bioengineering, 2016. 9(1): p. 73–84.

14. Polacheck, W.J., et al., A non-canonical Notch complex regulates adherens junctions and vascular barrier function. Nature, 2017. 552(7684): p. 258–262.

15. Miao, X., et al., Design, fabrication, and application of bioengineering vascular networks based on microfluidic strategies. Journal of Materials Chemistry B, 2025.

16. Yeo, M., et al., Synergistic coupling between 3D bioprinting and vascularization strategies. Biofabrication, 2023. 16(1).

17. Gold, K.A., et al., 3D Bioprinted Multicellular Vascular Models. Advanced Healthcare Materials, 2021. 10(21): p. 2101141.

18. Skylar-Scott, M.A., et al., Biomanufacturing of organ-specific tissues with high cellular density and embedded vascular channels. Science Advances, 2019. 5(9): p. eaaw2459.

19. Federici, A.S., et al., Muticomponent Melt-Electrowritten Vascular Graft to Mimic and Guide Regeneration of Small Diameter Blood Vessels. Advanced Functional Materials, 2024. 34(51): p. 2409883.

20. Ma, C., et al., Photoacoustic imaging of 3D-printed vascular networks. Biofabrication, 2022. 14.

21. Simińska-Stanny, J., et al., Borax - and tannic acid-based post-3D-printing treatment to tune the mechanical properties of scaffolds. Biomaterials Science, 2025. 13(13): p. 3689–3706.

22. Fazal, F., et al., Fabrication of a Compliant Vascular Graft Using Extrusion Printing and Electrospinning Technique. Advanced Materials Technologies, 2024. 9(23): p. 2400224.

23. Größbacher, G., et al., Volumetric Printing Across Melt Electrowritten Scaffolds Fabricates Multi-Material Living Constructs with Tunable Architecture and Mechanics. Advanced Materials, 2023. 35(32): p. 2300756.

24. Rizzo, R., et al., Optimized Photoclick (Bio)Resins for Fast Volumetric Bioprinting. Advanced Materials, 2021. 33(49): p. 2102900.

25. Kelly, B.E., et al., Volumetric additive manufacturing via tomographic reconstruction. Science, 2019. 363(6431): p. 1075–1079.

26. Riffe, M.B., et al., Multi-Material Volumetric Additive Manufacturing of Hydrogels using Gelatin as a Sacrificial Network and 3D Suspension Bath. Advanced Materials, 2024. 36(34): p. 2309026.

27. Bernal, P.N., et al., Volumetric Bioprinting of Complex Living-Tissue Constructs within Seconds. Advanced Materials, 2019. 31(42): p. 1904209.

28. Gehlen, J., et al., Tomographic volumetric bioprinting of heterocellular bone-like tissues in seconds. Acta Biomaterialia, 2023. 156: p. 49–60.

29. Krumins, E., et al., Glycerol-based sustainably sourced resin for volumetric printing. Green Chemistry, 2024. 26(3): p. 1345–1355.

30. Viola, M., et al., Thermal Shrinking of Biopolymeric Hydrogels for High Resolution 3D Printing of Kidney Tubules. Advanced Functional Materials, 2024. 34(46): p. 2406098.

31. McQueen, A. and C.M. Warboys, Mechanosignalling pathways that regulate endothelial barrier function. Current Opinion in Cell Biology, 2023. 84: p. 102213.

32. Peterson, L.W. and D. Artis, Intestinal epithelial cells: regulators of barrier function and immune homeostasis. Nature Reviews Immunology, 2014. 14(3): p. 141–153.

33. Wang, D., et al., Microfluidic bioprinting of tough hydrogel-based vascular conduits for functional blood vessels. Science Advances. 8(43): p. eabq6900.

34. Dessalles, C.A., et al., Integration of substrate- and flow-derived stresses in endothelial cell mechanobiology. Communications Biology, 2021. 4(1): p. 764.

35. Cunningham, K.S. and A.I. Gotlieb, The role of shear stress in the pathogenesis of atherosclerosis. Laboratory Investigation, 2005. 85(1): p. 9–23.

36. Fowler, M., et al., Guiding vascular infiltration through architected GelMA/PEGDA hydrogels: an in vivo study of channel diameter, length, and complexity. Biomater Sci, 2025. 13(11): p. 2951–2960.

37. Duan, J., et al., 3D Bioprinted GelMA/PEGDA Hybrid Scaffold for Establishing an In Vitro Model of Melanoma. J Microbiol Biotechnol, 2022. 32(4): p. 531–540.

38. Gao, J., et al., 3D-Printed GelMA/PEGDA/F127DA Scaffolds for Bone Regeneration. J Funct Biomater, 2023. 14(2).

39. Hall P, C.D., Wheeler T, Ostojich D, Aptecker J, Liashenko I, et al. x., A versatile and high-resolution hydrogel platform for volumetric additive manufacturing based on poly(ethylene glycol) diacrylate and alginate blends. ChemRxiv, 2024.

40. Hall, P., et al., A versatile and high-resolution hydrogel platform for volumetric additive manufacturing based on poly (ethylene glycol) diacrylate and alginate blends. 2024.

41. Rodríguez-Pombo, L., et al., Volumetric 3D printing for rapid production of medicines. Additive Manufacturing, 2022. 52: p. 102673.

42. Madrid-Wolff, J., et al., Controlling Light in Scattering Materials for Volumetric Additive Manufacturing. Advanced Science, 2022. 9(22): p. 2105144.

43. Xie, M., et al., Volumetric additive manufacturing of pristine silk-based (bio)inks. Nat Commun, 2023. 14(1): p. 210.

44. Bertana, V. and M. Periolatto, Volumetric 3D Printing, in High Resolution Manufacturing from 2D to 3D/4D Printing: Applications in Engineering and Medicine, S.L. Marasso and M. Cocuzza, Editors. 2022, Springer International Publishing: Cham. p. 131–151.

45. Falandt, M., et al., Spatial-Selective Volumetric 4D Printing and Single-Photon Grafting of Biomolecules within Centimeter-Scale Hydrogels via Tomographic Manufacturing. Adv Mater Technol, 2023. 8(15).

46. Young, A.T., O.C. White, and M.A. Daniele, Rheological Properties of Coordinated Physical Gelation and Chemical Crosslinking in Gelatin Methacryloyl (GelMA) Hydrogels. Macromol Biosci, 2020. 20(12): p. e2000183.

47. Ghosh, R.N., et al., An insight into synthesis, properties and applications of gelatin methacryloyl hydrogel for 3D bioprinting. Materials Advances, 2023. 4(22): p. 5496–5529.

48. Franck, A., Viscoelasticity and dynamic mechanical testing, T.I. Germany, Editor.

49. Shie, M.-Y., et al., Effects of Gelatin Methacrylate Bio-ink Concentration on Mechano-Physical Properties and Human Dermal Fibroblast Behavior. Polymers, 2020. 12(9): p. 1930.

50. Zhou, M., et al., Microbial transglutaminase induced controlled crosslinking of gelatin methacryloyl to tailor rheological properties for 3D printing. Biofabrication, 2019. 11: p. 025011.

51. Kavalli, T., Matériaux photopolymères avancés et systèmes de photo-amorçages pour l’impression 3D. 2019.

52. Hutson, C.B., et al., Synthesis and characterization of tunable poly(ethylene glycol): gelatin methacrylate composite hydrogels. Tissue Eng Part A, 2011. 17(13-14): p. 1713–23.

53. Beris, A.N., et al., Recent advances in blood rheology: a review. Soft Matter, 2021. 17(47): p. 10591–10613.

54. Goy, C.B., R.E. Chaile, and R.E. Madrid, Microfluidics and hydrogel: A powerful combination. Reactive and Functional Polymers, 2019. 145: p. 104314.

55. Weigel, N., et al., From microfluidics to hierarchical hydrogel materials. Current Opinion in Colloid & Interface Science, 2023. 64: p. 101673.

56. Xie, R., et al., Engineering of Hydrogel Materials with Perfusable Microchannels for Building Vascularized Tissues. Small, 2020. 16(15): p. 1902838.

57. Simińska-Stanny, J., et al., Advanced PEG-tyramine biomaterial ink for precision engineering of perfusable and flexible small-diameter vascular constructs via coaxial printing. Bioact Mater, 2024. 36: p. 168–184.

58. Ballermann, B.J., et al., Shear stress and the endothelium. Kidney International, 1998. 54: p. S100–S108.

59. Walji, N., S. Kheiri, and E. Young, Angiogenic Sprouting Dynamics Mediated by Endothelial-Fibroblast Interactions in Microfluidic Systems. Advanced Biology, 2021. 5.

60. Miranda Azpiazu, P., et al., A novel dynamic multicellular co-culture system for studying individual blood-brain barrier cell types in brain diseases and cytotoxicity testing. Scientific Reports, 2018. 8.

61. Galie, P.A., et al., Fluid shear stress threshold regulates angiogenic sprouting. Proceedings of the National Academy of Sciences, 2014. 111(22): p. 7968–7973.

62. Abello, J., et al., Peristaltic pumps adapted for laminar flow experiments enhance in vitro modeling of vascular cell behavior. Journal of Biological Chemistry, 2022. 298(10): p. 102404.

63. Morimoto, Y., et al., Microfluidic system for applying shear flow to endothelial cells on culture insert with collagen vitrigel membrane. Sensors and Actuators B: Chemical, 2021. 348: p. 130675.

64. Cuccia, N.L., et al., Pore-size dependence and slow relaxation of hydrogel friction on smooth surfaces. Proceedings of the National Academy of Sciences, 2020. 117(21): p. 11247–11256.

65. Simińska-Stanny, J., et al., Optimizing phenol-modified hyaluronic acid for designing shape-maintaining biofabricated hydrogel scaffolds in soft tissue engineering. Int J Biol Macromol, 2023. 244: p. 125201.

66. Madrid-Wolff, J., et al., A review of materials used in tomographic volumetric additive manufacturing. MRS Communications, 2023. 13(5): p. 764–785.

67. Simińska-Stanny, J., et al., Geometrical Designs in Volumetric Bioprinting to Study Cellular Behaviors in Engineered Constructs. bioRxiv, 2025: p. 2025.07.14.664683.

68. Delrot, P., Volumetric 3D printing for Life Science applications. 2024, Readily3D SA. p. 23.

69. Hapipi, N.M., et al., The Rheological Studies on Poly(vinyl) Alcohol-Based Hydrogel Magnetorheological Plastomer. Polymers, 2020. 12(10): p. 2332.

70. Falandt, M., et al., Hybrid supramolecular-covalent bioresin promotes cell migration and self-assembly in light-based volumetric bioprinted constructs. 2025.

71. Hospodiuk-Karwowski, M., et al., Dual-charge bacterial cellulose as a potential 3D printable material for soft tissue engineering. Composites Part B: Engineering, 2022. 231: p. 109598.

72. Komissarenko, D., et al., DLP 3D printing of high strength semi-translucent zirconia ceramics with relatively low-loaded UV-curable formulations. Ceramics International, 2023. 49(12): p. 21008–21016.

73. Simińska-Stanny, J., et al., Optimizing phenol-modified hyaluronic acid for designing shape-maintaining biofabricated hydrogel scaffolds in soft tissue engineering. International Journal of Biological Macromolecules, 2023. 244.

74. Shavandi, A., et al., Biomaterial ink based on bacterial polyglucuronic acid for tissue engineering applications. Next Materials, 2024. 4: p. 100181.

75. Schneider, S., et al., Peristaltic on-chip pump for tunable media circulation and whole blood perfusion in PDMS-free organ-on-chip and Organ-Disc systems. Lab on a Chip, 2021. 21(20): p. 3963–3978.

76. Cai, B., et al., One-step bioprinting of endothelialized, self-supporting arterial and venous networks. Biofabrication, 2025. 17(2).

