## Supplementary Information for "Controlling Flow Dynamics and Permeability in Perfusable Vascular Constructs Using Volumetric 3D Printing"

### *S1.1 Experimental section*

### *Formulation of biomaterial resins*

*Table S1 Different concentrations of methacrylate gelatin (GelMA), polyethylene glycol diacrylate (PEGDA), and lithium phenyl-2,4,6-trimethylbenzoylphosphinate (LAP) tested to adjust the resin for Vol3DP.*

| Resin formulations | GelMA (w/v %) | PEGDA (v/v %) | LAP (mg/mL) |
|---|---|---|---|
| **GelMA5/PEGDA10 LAP 1** | 5 | 10 | 1 |
| **GelMA5/PEGDA10 LAP 0,5** | 5 | 10 | 0.5 |
| **GelMA5/PEGDA10 LAP 0,3** | 5 | 10 | 0.3 |
| **GelMA10/PEGDA5 LAP 1** | 10 | 5 | 1 |
| **GelMA10/PEGDA5 LAP 0,5** | 10 | 5 | 0.5 |
| **GelMA10/PEGDA5 LAP 0,3** | 10 | 5 | 0.3 |
| **GelMA5/PEGDA5 LAP 1** | 5 | 5 | 1 |
| **GelMA5/PEGDA5 LAP 0,5** | 5 | 5 | 0.5 |
| **GelMA5/PEGDA5 LAP 0,3** | 5 | 5 | 0.3 |

### *S1.2 Results*

### *Volumetric printing investigations*

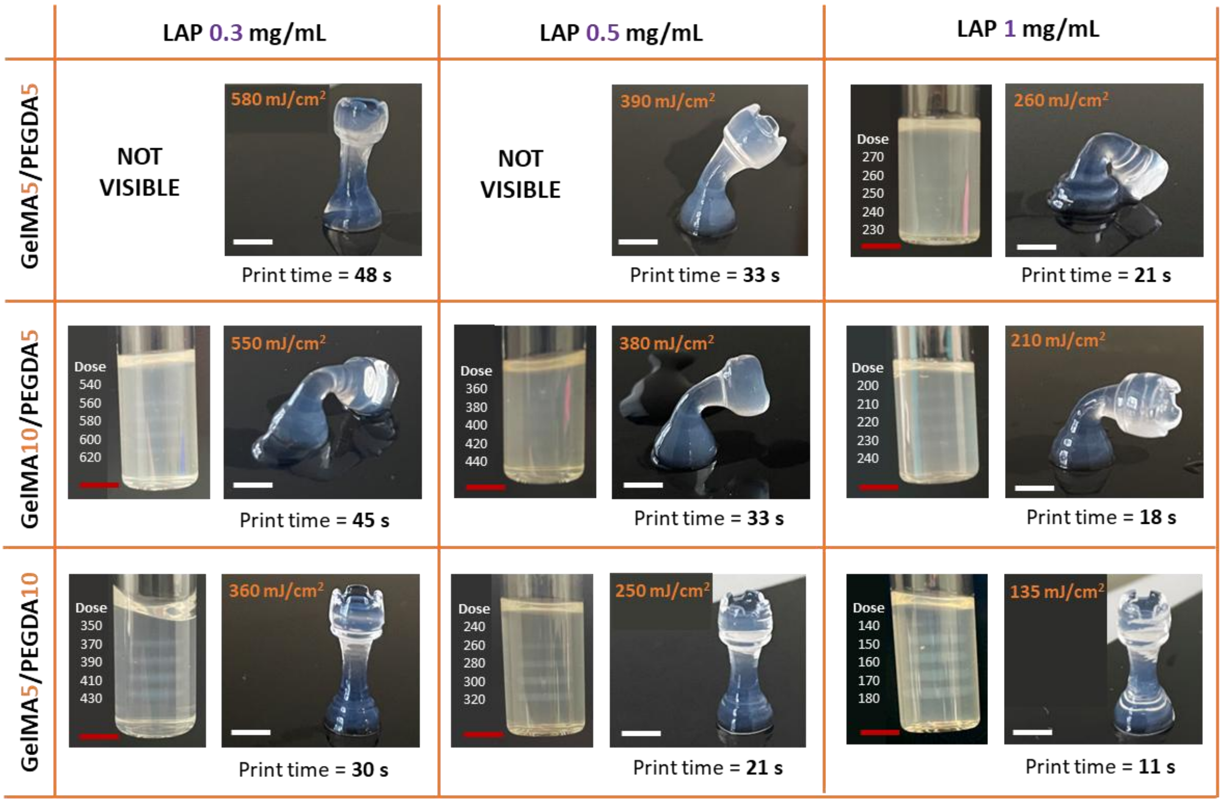


***Figure S1*** *Volumetric printability evaluation for various formulated resins with different photoinitiator (LAP) concentrations. Discs were printed to adjust the dose required for printing discs and chess pawns to demonstrate resin print resolution and shape-fidelity (n=3). The exposure time is obtained directly from the printer software, using a built-in algorithm to convert the minimum dose required into exposure time.*

### *Volumetric printing (Vol3DP) - Material and printing concerns*

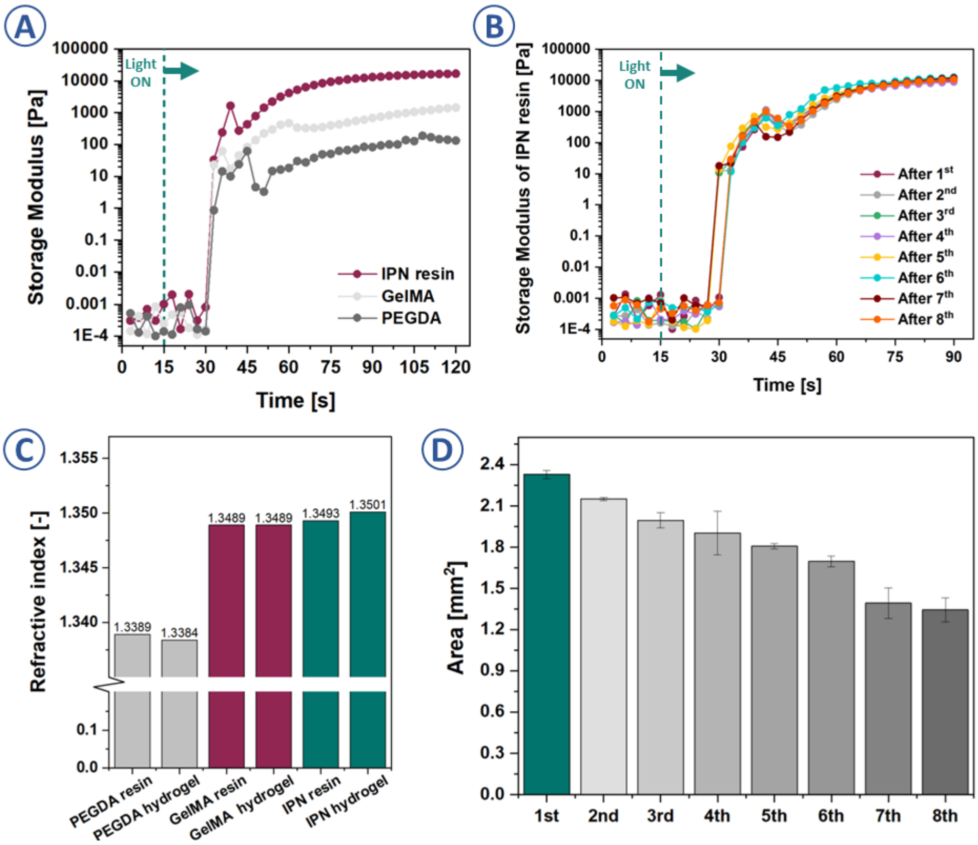


***Figure S2 A)*** *Photorheology tests for resin, GelMA and PEGDA (representative plots based on n=3), the dashed line represents the moment from which the 405 nm light was on;* ***B)*** *Photorheology tests for resin after every cooling and heating cycle (representative plots based on n=3), the dashed line represents the moment from which the 405 nm light was on;* ***C)*** *Refractive index measurements for resin and hydrogels based on GelMA, PEGDA and the resin used in this study;* ***D)*** *Area of the Vol3DP design after consecutive printing, n=3, mean ± SD., Perimeter of the Vol3DP design after consecutive printing, n=3, mean ± SD., results were significant at $p \leq 0.05$ only for $1^{st}$ vs. $7^{th}$ and $1^{st}$ vs. $8^{th}$ print, Kruskal-Wallis.*

### *Cells seeded within a multichannel Vol3DP hydrogel*

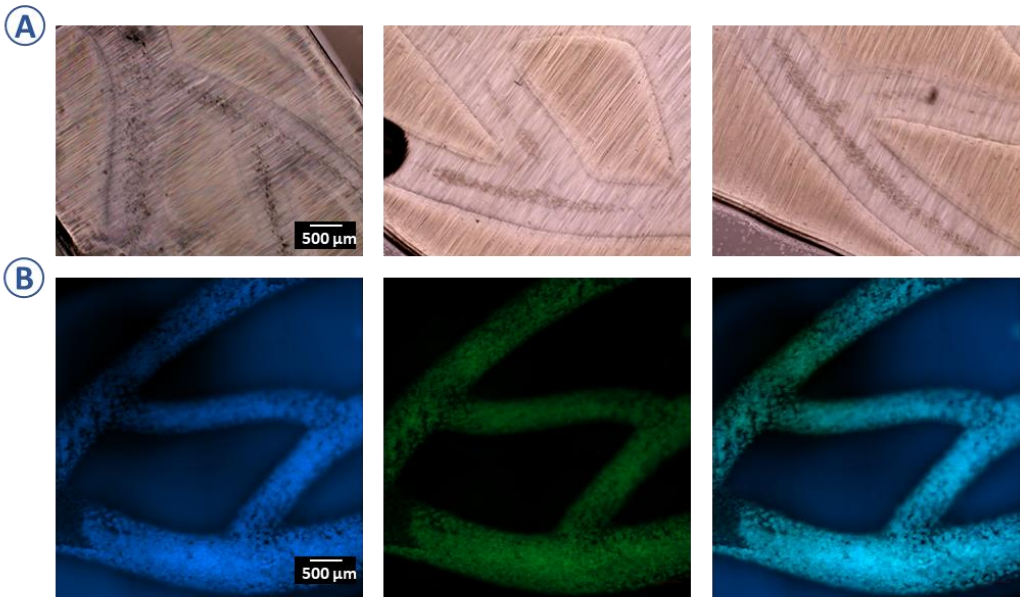


***Figure S3 A)*** *Vol3DP branched vascular model seeded cells after 8h of incubation;* ***B)*** *Images of the fluorescently labelled cells (day 7) lining the Vol3DP vascular model, the blue color depicts cell nuclei and green corresponds to the cytoskeleton.*

### *Permeation patterns within the Vol3DP hydrogel when perfused with dye*

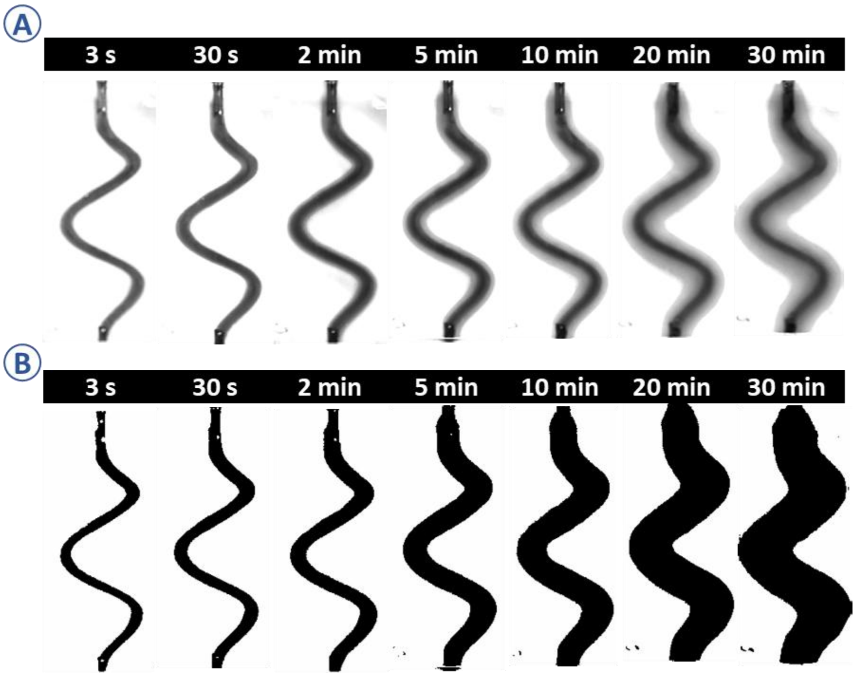


***Figure S4 A)*** *Grayscale images created in ImageJ based on permeation images taken at different time points;* ***B)*** *Binary images used to quantify the dyed area.*

### *CFD – simulated permeation patterns within the hydrogel*

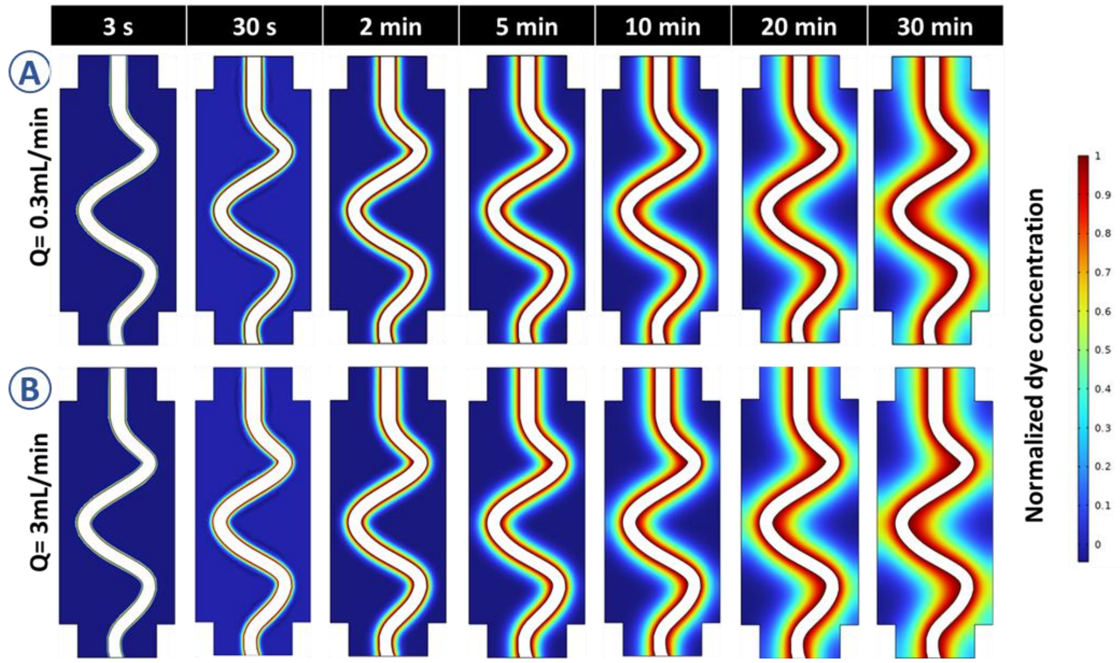


***Figure S5*** *Relative concentration of dye in the porous medium compared to that in the artificial blood vessel at different time points for fluid flows of **A)** 0.3 mL/min, **B)** 3 mL/min.*
